# Interplay of Spoonbill, Larp7 and survival motor neuron, in *Drosophila* model of Spinocerebellar Ataxia 8 non-coding RNA associated neurodegeneration

**DOI:** 10.64898/2025.11.30.691381

**Authors:** Rituparna Das, Pranjali Pandey, Bipin Tripathi, Bhawana Maurya, Ashim Mukherjee, Mousumi Mutsuddi

## Abstract

A deeper understanding of neurodegenerative disorders at the level of genetic or environmental risk factors as well as contributing genes, pathways, and networks suggests the presence of shared molecular mechanisms. Previously, we have reported that the KH domain of the Spoonbill protein alone can suppress non-coding Spinocerebellar Ataxia 8 (*SCA8*) associated neurodegeneration. In the current study, we have identified dLarp7 as a novel interacting partner of Spoonbill in *Drosophila*. Mammalian Larp7 is associated with a neurodevelopmental disorder, Alazami Syndrome (AS). In this study, we report that dLarp7 protein is recruited into the pathogenic *SCA8* RNA foci, which leads to depletion of its downstream target, *7SKsnRNA*. Soaking away of Larp7 into toxic RNA foci results in its depletion from the physiological pool resulting in destabilization and depletion of 7SKsnRNP. Hence, increasing the dose of dLarp7 suppressed *SCA8* associated neurodegeneration, by restoring the physiological levels of dLarp7 and 7SKsnRNP. In addition, it was observed that dLarp7 interacts with the somatic motor neuron (SMN) protein which is associated with spinal muscular atrophy (SMA). This observation led us to explore the interaction of *Drosophila SMN1* orthologue with pathogenic *SCA8* associated neurodegeneration. Intriguingly, SMN protein modulated molecular neuropathogenesis associated with *SCA8*. The presence of orthologues of *Drosophila* RNA binding proteins, dLarp7 and SMN, with mammalian counterparts underlines the translational importance of our findings. Our study also hints at the shared molecular mechanisms that underlies multiple neurodegenerative diseases. A novel association of *Drosophila* homologs of AS-linked Larp7 and SMA-causing SMN1 with *SCA8* associated neurodegeneration suggests an overlap of molecular threads underlying the pathogenesis of neurodegenerative disorders.

## INTRODUCTION

Repeat expansion mutations in non-coding genes cause RNA repeat expansion neurodegenerative disorders (RENDs) (Das et al. 2019). RENDs include Myotonic Dystrophy 1 and 2 (DM1, 2), Fragile X-Associated Tremor/Ataxia Syndrome (FXTAS), Huntington’s disease-like 2 (HDL2), *C9orf72*-associated Amyotrophic lateral sclerosis/frontotemporal dementia (*C9orf72*ALS/FTD), and Spinocerebellar Ataxia 8, 10, 31, 36 (Todd & Paulson, 2010); Rodriguez and Todd 2019). Many of the above diseases have been modelled in *Drosophila,* because of its highly tractable molecular genetic tools, conserved signaling pathways along with striking homology with human disease-causing genes (Das et al. 2019).

We have earlier generated a *Drosophila* spinocerebellar ataxia 8 (*SCA8*) neurodegenerative disorder model (Mutsuddi et al, 2004). The expanded CTG repeat in the 5’UTR of *ATXN8OS* is associated with *SCA8*. *SCA8* afflicted patients have gait instability, mild aspiration, unclear speech, progressive inability to walk along with development of cerebellar degeneration. *Drosophila SCA8* model that ectopically express pathogenic *CTG(112)* repeats in the photoreceptor neurons results in severe photoreceptor disorganization that worsens over time (Mutsuddi et al, 2004). Our disease model mimics human *SCA8* neurodegenerative disease phenotypes like formation of pathogenic RNA foci and progressive neurodegeneration. Drosophila *SCA8* model was used to screen for novel modifiers and Spoonbill (Spoon) was identified as a suppresser of the neurodegenerative disease phenotype. Previously we have reported that the KH domain of Spoon physically interacts and depletes the expanded pathogenic *SCA8* RNA (Tripathi et al., 2016). Spoon, which is an RNA binding protein, interacts with the neuronal cell-fate determining factor Prospero (Tripathi et al., 2017) and modulates JNK signaling (Das et al. 2023). Overexpressing Spoon KH domain alone suppressed *SCA8* neurodegenerative phenotype. Thus, KH domain of Spoon in future can be targeted as a therapeutic endpoint for *SCA8*-related neurodegeneration (Mutsuddi & Rebay, 2005; Wang & Chang, 2011). The repeat expanded toxic *SCA8* transcripts form an unusual RNA-protein complex with Spoonbill, suggesting its recruitment into the pathogenic molecular sink. Proteomic analysis of Spoon was done to identify its interacting partners that modulates *SCA8* associated neurodegeneration. From this screen, we identified *Drosophila* RNA binding protein dLarp7 as a physical interactor of Spoon.

Larp7 is a vital RNA-binding protein with two RNA recognition motifs and one RNA binding domain. Mammalian Larp7 was first identified as an oligo-U binding protein (Markert et al., 2008; Wang & Chang, 2011)) and acts as a chaperone for *7SKsnRNA* (Brogie & Price, 2017; Eichhorn et al., 2018; Krueger et al., 2008). 7SKsnRNP constitutes of Larp7, HEXIM, hnRNPA1, and MEPCE proteins. It retains P-TEFb in an inhibitory complex (Markert et al. 2008; Krueger et al., 2008). Active P-TEFb, consisting of Cdk9, CyclinT1 and 2, phosphorylates the C-terminal domain of the poised RNA Polymerase II and is involved in transcription elongation (Ni et al., 2008).

Interestingly, Alazami Syndrome (AS; MIM 615071), a rare autosomal recessive neurodevelopmental disorder is associated with Larp7 loss of function mutation (Alazami et al., 2012)(Wojcik et al., 2019; S. Das et al., 2021). Alazami patients are characterized by dysmorphic facial features, developmental delay along with intellectual disability. AS patients with mutations in Larp7 have short lymphocyte telomeres (Holohan et al., 2016) and also impairs RNA splicing (Wang et al. 2020; Hasler et al. 2020). Mammalian Larp7 regulates telomerase activity and splicing, genome stability in cancer cells as well as mitochondrial biogenesis (Yu et al., 2021; Zhang et al., 2020). The role of Larp7 in neurodevelopmental disorder further motivated us to explore the function of its *Drosophila* homolog in the *SCA8* non-coding trinucleotide repeat associated neurodegeneration. *Drosophila* Larp7 (dLarp7) shares 32% identity and domain organization with mammalian Larp7. Similar to that of the mammalian 7SKsnRNP, dLarp7 is vital for the stability of *Drosophila 7SKsnRNA* and 7SKsnRNP (Nguyen et al., 2012).

In this study we report that dLarp7 modulates Spoonbill dependent pathogenic *SCA8* associated neurodegeneration in the developing photoreceptor neurons. Overexpression of dLarp7 leads to the depletion of pathogenic *SCA8* RNA foci, which is the hallmark of the disease along with reduction in pathogenic *SCA8* transcript levels. Further, RNA *in situ* hybridization (RISH) along with immunostaining and RNA immunoprecipitation revealed that dLarp7 is recruited into the pathogenic *SCA8* RNA foci. This probably results in destabilization of 7SKsnRNP and contributes to *SCA8* associated neuropathology. Identification of dLarp7 as a potent suppressor of *SCA8* associated neurodegeneration implicates shared mechanisms underlying neuronal disorders.

In mice motor neurons, spliceosome biogenesis is precisely regulated and coupled to transcriptional demand through cell-specific RNA-binding proteins, chromatin modifications, and the physical interaction between the transcription and splicing machineries. Interestingly, it has been reported that 7SK core component Larp7, interacts with Somatic motor neuron (SMN) protein to regulate spliceosome biogenesis in accordance with the transcriptional demand in mice motor neuron cells (Ji et al., 2021). SMN protein is vital for organization and stability of spliceosomal UsnRNPs. SMN deficiency causes spinal muscular atrophy (SMA), which is an autosomal recessive neurodegeneration affecting spinal motor neurons. We show that *Drosophila* SMN interacts with dLarp7 and largely colocalizes with Spoonbill in photoreceptor neurons. Intriguingly, *Smn* significantly modulated *SCA8* associated neurodegeneration which also regulated toxic RNA foci formation. This further consolidated the possibility of the presence of a shared molecular machanism underlying neuronal diseases.

The orthologous nature of *Drosophila* dLarp7 and SMN proteins with their human counterparts highlights the therapeutic relevance of our study. Identification of *Drosophila* homologs of well-established disease associated genes like Larp7 and SMN, in *SCA8* associated neurodegeneration, opens up a possibility of the presence of shared mechanisms underlying neuronal diseases. We have previously identified RNA binding proteins (RBPs) like muscleblind (mbl/mbnl1) as suppressors for spinocerebellar ataxia 8 (*SCA8*). mbnl has also been implicated in myotonic dystrophy 1 (DM1), and Amyotropic Lateral Sclerosis (ALS) suggesting a shared process underlying these disorders. (Ji et al., 2021). We propose that loss of such vital RNA binding proteins either due to mutation or unusual recruitment into pathogenic RNA foci destabilizes the critical balance of transcription and splicing thereby contributing to neuropathology. Perhaps, such a common molecular mechanism underlies multiple neurodegenerative disorders. Further, research into the structural aspect of this RNA-protein interaction and their targets, in disease models and patients will help in fortifying our findings. Identification of specific molecular changes for a range of neurodegenerative disorders early in the progression of the disease will provide shared biomarkers and therapeutic targets. Hence, understanding the shared molecular mechanisms underlying multiple neurodegenerative disorders can lead to the identification of common therapeutic targets for the management of multiple neurodegenerative disorders and provide robust biomolecular markers for early diagnosis.

## MATERIAL AND METHODS

### Fly stocks and rearing

Fly stocks were maintained on standard cornmeal/yeast/molasses/agar medium at 25°C. Genetic interaction studies with *UAS SCA8(CTG112)* and *SCA8(CT9*) were performed at 28°C. *Oregon^R^, w^1118^,* and *w; GMR-GAL4/+;+/+* was used as control for experiments. *w; +/+; UAS-SCA8(CTG9) /TM6BTb* and *w; +/+; UAS-SCA8(CTG112)/TM6BTb* were generated previously (Mutsuddi et al. 2004). *UAS-HA-spoonbill* was generated in our lab (Tripathi et al. 2016), while other alleles of spoonbill*UAS-spoon^EP1400^* (BL 11236) and *spoonbill^KG02745^* were obtained from Bloomington Stock Center. BL 21200, BL 32898, BL 31762, and BL 41932 were also obtained from Bloomington Stock Center.

### Generation of HA tagged transgenic lines

LD09531, BDGP Gold cDNAs clone for *dLarp7* was obtained from the *Drosophila* Genomics Research Center and was utilized as the template for cloning full length CDS of *dLarp7* with HA-tag sequence at the 5’ end, into the *pUAST* vector employing approach described previously (Tripathi et al. 2016). The recombinant constructs were sequence verified and microinjected into pre-blastoderm embryos of *w^1118^* flies following standard germ line transformation procedures for *Drosophila.* The transgenic lines were verified for dLarp7 overexpression (Figure S1).

### Generation of dLarp7 RNAi transgenic lines

Sub-cloning was performed in *pUAST-attB* vector for generating two RNAi lines of dLarp7. The recombinant plasmids sequences were verified and sent for microinjection and development of transformant lines. The transgenic lines obtained, were verified for depletion of dLarp7 protein and transcripts (Figure S2).

### Generation of anti-dLarp7 antibody

Antibodies against dLarp7 protein was generated by subcloning a unique short 150bp N-terminal sequence of dLarp7 in the *pGEX-4T1* vector as described previously (Tripathi et al. 2017). The IPTG-induced recombinant peptide was pulled down and eluted; and was used for generation of antibody in rats. The antisera from all the 4 rats were tested for specificity by immunostaining against dLarp7(Figure S2).

### Imaging of Adult Eye Tissues

Age matched 2-3 days post-eclosion flies with the desired genetic combination were collected and scored for their phenotypes. The etherized flies were mounted on a bridged slide for bright field microphotography. For representation purpose, only females were reported in the panels. Similarly, eye imprints were prepared using transparent nail paint and documented using Nikon Eclipse Ni.

### Immunocytochemistry and microscopy

*Drosophila* third instar larval tissues were dissected out for the immunostaining and was performed as described previously (Das et al., 2023). Following primary antibodies were used in this study: Mouse anti-HA (1:100, Sigma), mouse anti-22C10 (1:300, DSHB), Alexa555-, or Alexa405-conjugated secondary antibodies against mouse (1:200) and rat (1:200) were used to detect the primary antibodies. Nuclear counterstaining was done using DAPI and mounted in DABCO (Sigma). The slides were observed under Carl Zeiss LSM 780 laser scanning confocal microscope and images were processed with Adobe Photoshop7. The intensity profiling was done using Image J and Graphpad Prism software 5.0.

### RNA purification and Real-Time PCR

Adult fly heads from suitable genetic combinations were collected for RNA isolation using method described previously (Das et al., 2023). 2µg DNaseI (Roche) treated RNA was reverse transcribed using M-MuLV Reverse Transcriptase (NEB) following the manufacturer’s instructions. Changes in *SCA8*(CTG112) and *7SKsnRNA* transcript levels were measured using real-time PCR (QuantStudios5, Applied Biosystems) with 2X of Maxima SYBR-Green master mix (Thermo). The RT-PCR analysis was performed using the Livak method, and the fold changes were computed using 2^−ΔΔCT^ normalised to Rps17 (Schmittgen & Livak, 2008).

### Acridine orange staining

Eye-antennal discs were dissected from the third instar larvae in 1×PBS. Tissues were incubated immediately in 1 μg/ml acridine orange (Sigma) for 1.30 minutes. After 2 washes of 1 minute each, tissues were mounted gently in PBS and immediately observed under the fluorescence microscope (Nikon Eclipse Ni).

### Fluorescent RNA: RNA in situ hybridization

DIG-labeled *SCA8* and *hsrɷ93D* specific riboprobes were generated by *in vitro* transcription

The pBSK(-)*SCA8* plasmid DNA clone was linearized with *NotI* and *XhoI* for generating DIG-labeled *SCA8* specific antisense and sense riboprobes, respectively. 1μg of this linearised plasmid was used as a template for preparing probes with uridines labeled with digoxigenin using manufacturer’s protocol (Roche). The labeled RNA was precipitated by adding 10μg of yeast tRNA (Sigma) and 2.5 volumes of chilled ethanol at −20 °C overnight. The labeled RNA was then collected by centrifuging at 16,000 g for 15 minutes at 4 °C. The pellet was washed in 80% ethanol, air dried and dissolved in DEPC treated MQ.

RNA:RNA *in situ* hybridization (RISH) was carried out in intact tissues essentially as described earlier (Lehmann & Tautz, 1994)with slight modifications. Hybridization was carried out in hybridization buffer A with 100 ng *SCA8* antisense riboprobe at 50 °C for 12–16 h. As a positive control *hsrω* antisense riboprobe was hybridized for 12–16 h at 50 °C. Sense probes against *SCA8* was utilized as negative control (Figure S3). Probes were detected with anti-DIG-Rhodamine conjugated antibody (1:200, Roche). and counterstaining of the nucleus was done using DAPI. The RNA foci were counted with the help of LSM510 Meta software.

### Immunofluorescence with RNA *in situ* hybridization

For combined *in situ* detection of protein and RNA, immunofluorescence was performed prior to RNA hybridization in eye-antennal discs. Immunofluorescence was performed in RNase-free condition, along with excess heparin and tRNA for minimizing antibody non-specificity and protecting the RNAs for subsequent detection. The tissues were fixed twice, once after dissection and again after immunofluorescence, to crosslink the secondary antibody and retain fluorescence after RNA hybridization. Immunofluorescence with RNA *in situ* hybridization was performed as described earlier, with some modifications (Toledano et al., 2012). Immunostaining against HA-dLarp7 was followed by RISH against *SCA8*(CTG112) transcripts for detecting localization of the RNA and protein.

### Immunoprecipitation of RNA-protein complex

Immunoprecipitation of RNA-protein complex was performed as described previously (Tripathi et al. 2016). Anti-HA conjugated agarose beads were equilibrated and added to the lysates with HA-dLarp7 along with 1μg of *in vitro* transcribed *SCA8*(CTG112) RNA. Equal amounts of protein was used for western blotting β-tubulin and HA-dLarp7. RNA was eluted from the beads by vortexing the suspension in 100μl of phenol–chloroform–isoamyl alcohol followed by a brief centrifugation for 1 minute at 4°C to collect the RNAs bound with the beads. Yeast tRNA, sodium acetate and ethanol were added to precipitate the RNA from the aqueous phase. The RNA obtained from the pulled down product was then used to synthesize the first-strand cDNA synthesis using M-MuLV Reverse Transcriptase (New England Biolabs), followed by RT-PCR for *7SKsnRNA* (Figure S4.) and *SCA8*(CTG112).

### Statistical analysis

Intensity profiles were generated using Image J and represented in terms of intensity per unit area. Statistical analysis of genetic interactions, intensity profiles and climbing analysis was done using the GraphPad Prism 5 software. The error bars represent the standard error of the mean value of the replicated experiments. To determine the significance of our data ANOVA with Tukey’s multiple comparison post-test or Unpaired *t-*test were employed for the different genotypes. *p*-value <0.05 was accepted as statistically significant.

## RESULTS

### 1. Identification of dLarp7 as an interacting partner of Spoonbill

Proteomic analysis was carried out to identify the novel components of Spoon-*SCA8* RNA-protein complex. Proteins interacting with Spoon were pulled down by HA-conjugated beads incubated in protein lysates comprising of overexpressed HA tagged Spoon and pathogenic *SCA8* RNA. The interacting proteins were identified using MALDI-MS/MS. This led to the identification of five unique peptides that corresponded to *Drosophila* Larp7 (dLarp7). Thus, MADLI-MS/MS based proteomic screening identified dLarp7 as an interacting partner of Spoon (Figure 1. A).

**Figure 1.**
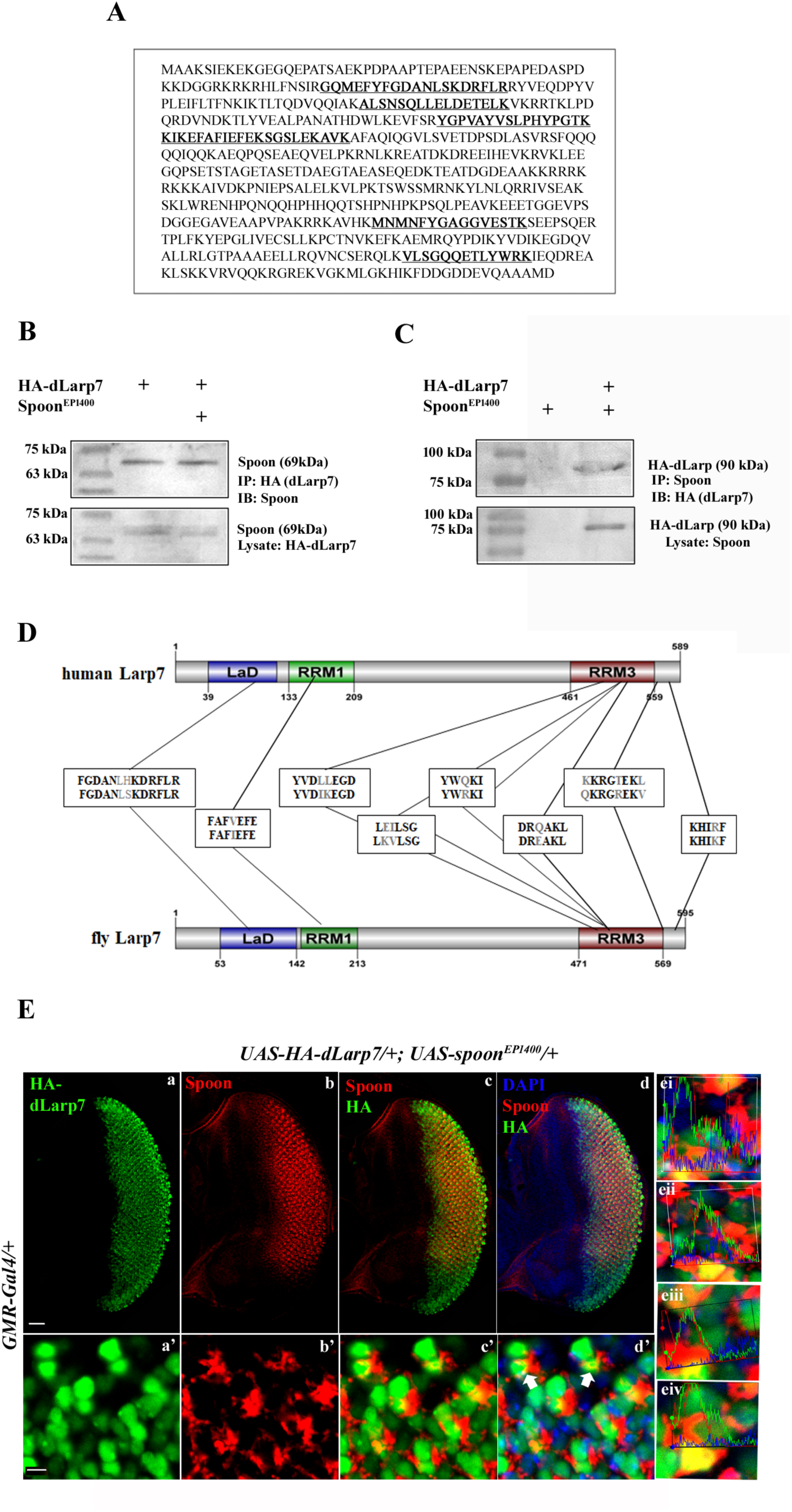
dLarp7 interacts with Spoonbill. (A) Mass spectrometric analysis identified peptide sequences that correspond to *Drosophila* dLarp7 protein. The Spoonbill protein complex was pulled down in lysates with overexpressed *spoon* with and without *SCA8*, and was run on SDS-PAGE. Mass spectrometry of a unique band using MALDI-MS/MS analysis identified six peptides unique to *Drosophila* dLarp7. (B, C) Co-immunoprecipitation of HA-dLarp7 and Spoonbill. Both the proteins were overexpressed in *Drosophila* photoreceptors and co-immunoprecipitated using anti-HA conjugated beads (B) or anti-Spoon antibody (C). Western blot with anti-Spoon (B) or anti-HA (C) antibody revealed presence of dLarp7 and Spoonbill in the same protein complex. Only single band was obtained in IP, whereas input lysates of HA-dLarp7 show two bands for Spoonbill protein (lower panel, Tripathi et al. 2017). (D) *Drosophila* dLarp7 has identical domain structure as human dLarp7, with multiple conserved motifs. Clustal Omega by EMBL-EBI was used for multiple sequence alignment to reveal nearly 32% identity between the two proteins. (E) Colocalization of dLarp7 and Spoonbill in *Drosophila* eye antennal. HA-tagged dLarp7 and Spoonbill were co-expressed in *Drosophila* eye antennal discs. dLarp7 was detected using anti-HA antibody (a, a’) and Spoonbill was visualized using anti-Spoonbill antibody (b, b’). The merged panel (c, c’) showed dLarp7 and Spoonbill colocalized partially, prevalent in the perinuclear region (d, d’). Intensity peaks validate overlapping signals from both channels indicating a significant colocalization of the two proteins (ei-iv). Scale bars, 50µm (a), 5 µm (a’).

In order to investigate the cooperative role of dLarp7 and Spoon for mediating suppression of pathogenic *SCA8* associated neurodegeneration, transgenic lines expressing complete coding sequence of dLarp7 with an HA tag at the 5’ end under UAS regulator were generated. Three transgenic lines were obtained which were then verified for ectopic dLarp7 expression using RT-PCR and Immunostaining (Figure S1).

Co-immunoprecipitation was carried out to confirm the physical interaction between dLarp7 and Spoonbill protein. Spoon protein was pulled down with HA-tagged dLarp7 in lysates containing both the proteins which were ectopically driven by *GMR-GAL4* driver (Figure 1. B, C). *spoon* encodes two protein isoforms which can be seen as two very close bands around 69kDa. However, the lane containing the co-immunoprecipitated proteins (Figure 1. B) displayed a single protein band of Spoon, suggesting that dLarp7 interacts with only one of the isoforms of Spoon. Thus, the proteomic analysis implicated that Spoonbill and dLarp7 are part of the same protein complex and this was verified and confirmed. *Drosophila* dLarp7 includes domains which are identical in organization with that of the human dLarp7 (Figure 1. D). dLarp7 has one La domain and two RNA recognition motifs that harbor conserved sequences with that of the human Larp7 (Figure 1. D).

This was followed up co-localization studies using immunocytochemistry. Immunostaining using anti-HA for detecting HA-dLarp7 revealed it to be largely nuclear in subcellular localization (Figure 1. E, Figure S1) that paralleled earlier reports (Nguyen et al. 2012). However, in the developing eye-antennal discs cells, some of the cells revealed cytoplasmic localization, mostly clustered around the morphogenetic furrow, suggesting a transient cytoplasmic localization during development (Figure 1. E, a, a’, Figure S1 D). Spoon protein on the other hand is localized outside the nucleus (Figure 1. E, b, b’) (Zhang et al. 2016). Colocalization analysis using anti-Spoon against Spoonbill and anti-HA antibodies dLarp7 respectively in the eye-antennal discs of the third instar larvae of *Drosophila* suggested that the two proteins colocalize partially and transiently in the cytoplasm (Figure 1. E, c, c’ d’). Overlapping graphs of red and green channels in intensity profiling of such points further supported our observation (Figure 1. E, ei-iv). Taken together, our observations indicate that dLarp7 is a novel interacting partner of Spoonbill in *Drosophila*.

### 2. dLarp7 suppresses *SCA8* associated phenotype in *Drosophila*

Ectopic expression of pathogenic *SCA8*(CTG112) in the developing photoreceptor of *Drosophila* resulted in a disorganized fused ommatidial array resulting in progressive degeneration of the photoreceptor neurons (Figure 2. A b, b’) (Mutsuddi and Rebay 2005; Mutsuddi et al. 2004). We have earlier identified *spoon* as a suppressor of pathogenic *SCA8* associated neurodegeneration. In order to unravel the molecular mechanism underlying *spoon* mediated suppression of pathogenic *SCA8*, role of its novel interacting partner dLarp7 was examined in *SCA8* associated neurodegeneration.

**Figure 2.**
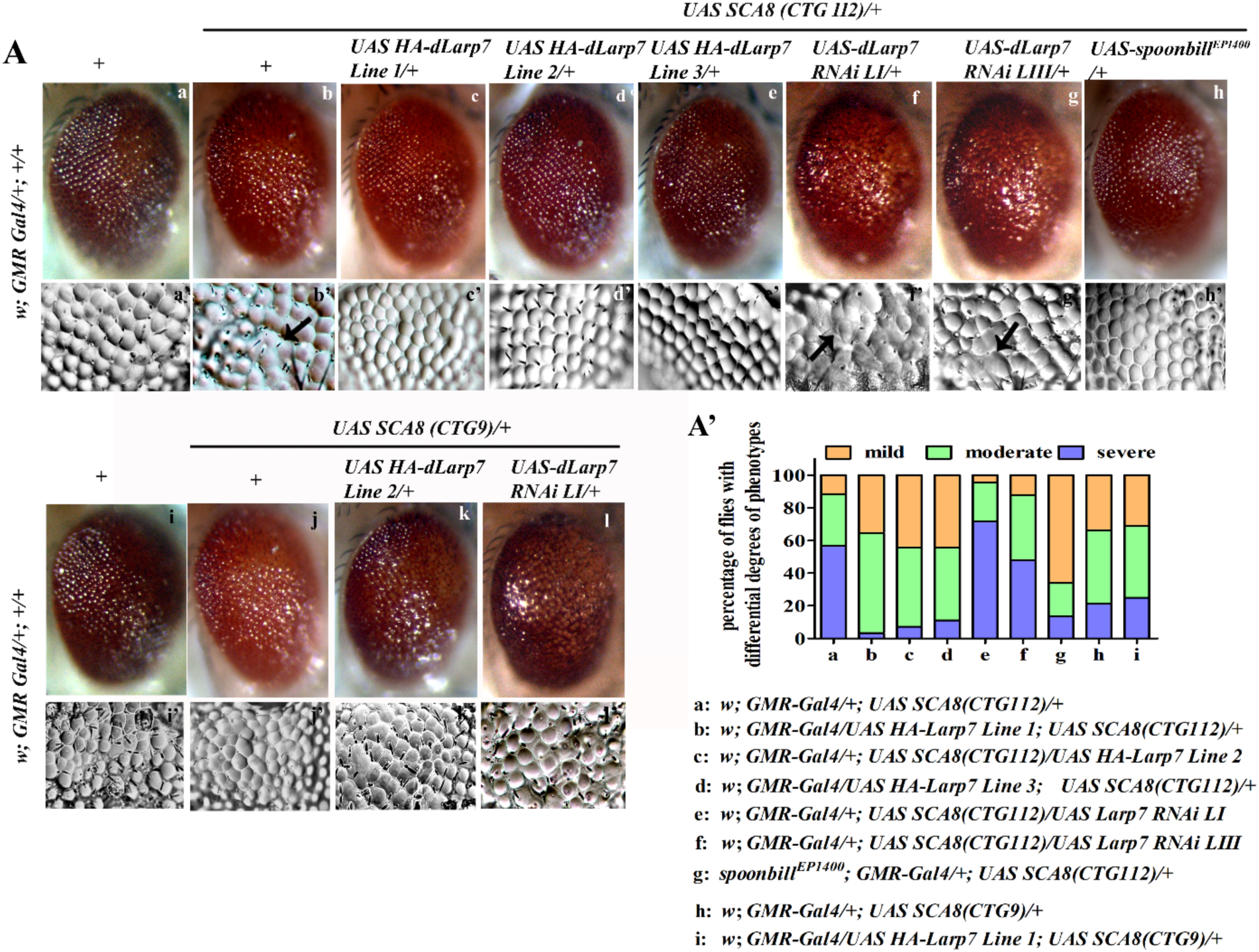
dLarp7 results in target specific modulation of *SCA8*(CTG112) associated neurodegeneration. dLarp7 overexpression alleles and RNAi lines modulate *SCA8(CTG112)* associated eye phenotype. *GMR-Gal4* driven ectopic expression of *UAS SCA8(CTG112)* show disorganized array (b) and fused clusters of ommatidia (b’) as seen using bright field micrography and nail paint imprinting compared to driver control (a, a’). Overexpression of *Drosophila dLarp7* with this sensitized background of ectopic *UAS SCA8(CTG112)* rescued the rough eye phenotype in adults (c, c’-e, e’). Combining *dLarp7 RNAi LI* or *LIII* with *UAS SCA8(CTG112)* resulted in enhanced neurodegenerative phenotype (f, f’ – g, g’). Expressing non-pathogenic *UAS SCA8(CTG9)* in *Drosophila* photoreceptors result in only a moderate eye roughening (j,j’ vs i, i’). Genetically combining overexpression allele of dLarp7 (k, k’) or dLarp7 RNAi (l, l’) with *UAS SCA8(CTG9)* do not modulate the eye phenotype. (A’) Quantitative assessment based on percentage phenotypes distributed based on severity of the phenotype revealed a pronounced rescue of severe eye phenotype by dLarp7 overexpression lines. Similarly, severity of eye phenotype was seen to be massively enhanced in combinations with dLarp7 RNAi LI and *UAS SCA8(CTG112)*, along with significant increase in the proportion of flies demonstrating severe eye phenotype.

Transformants expressing full length *UAS HA-dLarp7* were generated. The severe rough and disorganized ommatidial phenotype associated with pathogenic *SCA8*(CTG112) repeats was used as a sensitized background to assess the role of dLarp7 in modulating neurodegeneration. Rough eye phenotype associated with *UAS SCA8*(CTG112) (Figure 2. A b, b’ vs a, a’) was suppressed in combination with all the three transgenic lines expressing *UAS HA-dLarp7* (Figure 3. A c, c’, d, d’ e, e’ vs b, b’). All the three transgenic lines suppressed the *SCA8*(CTG112) induced eye roughening (Figure 3. A c, d, e vs b) as well as fused ommatidia (Figure 3. A c’, d’, e’ vs b’). On similar lines, transgenic dLarp7 RNAi lines with depleted dLarp7 enhanced *SCA8*(CTG112) pathogenic phenotype (Figure 3. A f, g vs b). In our control experiments, the mild eye roughness associated with non-pathogenic *SCA8*(CTG9) (Figure 2. A j, j’ vs i, i’) underwent no modification in combination with dLarp7 over expression or depletion of dLarp7 (Figure 2. A k, k’, l, l’ vs j, j’). Hence, these observations indicates that dLarp7 suppresses neurodegenerative phenotype associated with trinucleotide repeat associated toxic *SCA8*(CTG112) transcripts in a target specific manner, and this paralleled suppression by Spoon. (Figure 2. A h, h’)

**Figure 3.**
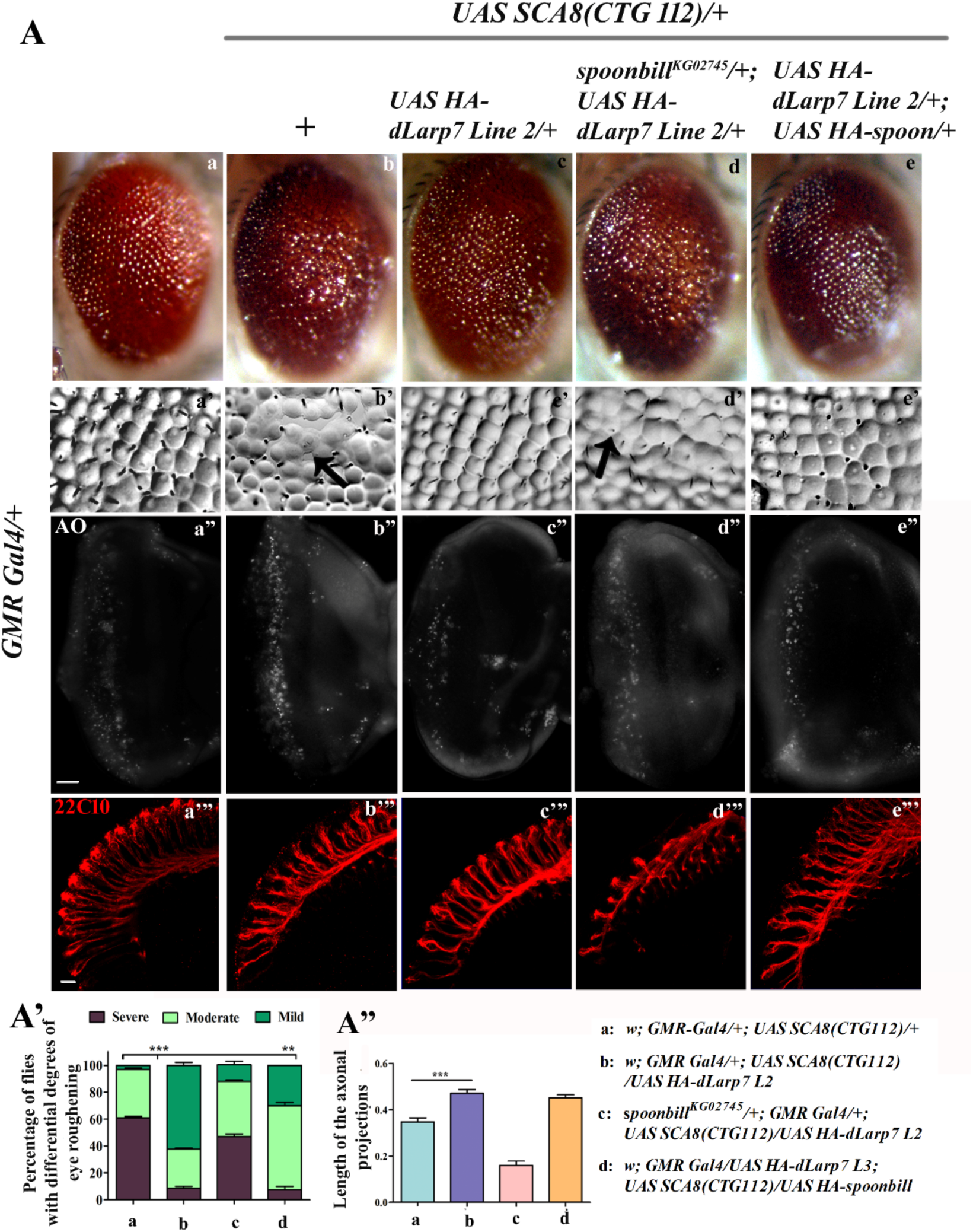
Overexpression of *dLarp7* suppresses pathogenic *SCA8*(CTG112) associated neurodegeneration in *Drosophila.* (A) *GMR-Gal4* driven ectopic expression of *UAS SCA8(CTG112)* show disorganized array (b) and fused clusters of ommatidia (b’) as seen using bright field microscopy and nail paint imprinting. Acridine Orange marks cells death in eye-antennal discs due to ectopic *UAS SCA8(CTG112)* expression (b”). 22C10 staining showed disturbed neuronal morphology and axonal projection in larval eye-antennal discs of *GMR-Gal4* driven *UAS SCA8(CTG112)* (b”’). Overexpression of *Drosophila dLarp7* in this sensitized background of ectopic *UAS SCA8(CTG112)* rescued the rough eye phenotype in adults (c, c’). *UAS-HA-dLarp7* rescued axonal morphology (c”’) and cell death in *Drosophila* larval eye-antennal discs (c”). Depletion of Spoonbill in the background of *GMR-Gal4* driven *UAS SCA8(CTG112)* and *dLarp7* interferes with the dLarp7 induced rescue of *SCA8* associated neurodegenerative phenotypes in *Drosophila* (d, d’, d”, d”’). Co-expression of *UAS-HA-spoonbill* along with *GMR-Gal4* driven *UAS SCA8(CTG112)* and *UAS-HA-dLarp7* results in no additional rescue over *dLarp7* overexpression (e, e’, e”, e”’). (A’) Quantification of the genetic interactions was done by calculating the percentage of flies showing the kinds of adult eye phenotypes. Percentage of flies showing severe eye phenotype reduced drastically on over expressing *UAS-HA-dLarp7* in *UAS SCA8(CTG112)* background. The modulation of percentage of flies showing severe and mild phenotypes was very significant between b and a. (A”) Quantitative assessment of the length of axonal projection also revealed a significant rescue by *UAS-HA-dLarp7.* To calculate statistical significance of the data Bonferroni pot test was added after two-way ANOVA analysis for the genetic interactions. Bars represent average ± SEM for two independent experiments. p values < 0.001***. Scale bars, 50 µm (A a”), 10µm (A a”’).

Quantification of the phenotypes obtained via genetic interactions were done by segregating the progeny into severe, moderate and mild phenotypes. Quantification revealed reduction in the percentages of flies with severe phenotype when Line 1 *UAS-HA-dLarp7* was used, this was followed closely by Line2. Hence, unless mentioned, in further experiments only Line 1 was used. The rescue seen by all three lines of *UAS HA-dLarp7* was very similar to that by overexpression of Spoon (Figure 2. A’). Enhancement of the neurodegenerative phenotypes associated with pathogenic *SCA8* non-coding RNA in combination with *UAS-dLarp7 RNAi LI* was found to be more pronounced than with *UAS-dLarp7 RNAi LIII.* Thus, these genetic interactions suggested that *dLarp7* is a potent suppressor of pathogenic *SCA8* associated neurodegeneration, which resonated Spoon mediated suppression. Again, no modulation in the eye phenotype of non-pathogenic *SCA8*(CTG9) was observed, clearly depicting that the loss in dLarp7 enhanced the phenotype is specifically associated with the pathogenic *SCA8* transcripts.

### 3. Suppression of pathogenic *SCA8* associated neurodegeneration by ectopic dLarp7 is mediated via Spoon

In order to better understand the role of dLarp7 in the suppression of *SCA8*(CTG112) associated neurodegeneration, the combinatorial role of Spoon and dLarp7 was also assessed. The pathogenicity of *SCA8*(CTG112) was analyzed in the adult photoreceptors as well as in eye-antennal discs from the III^rd^ instar larvae. dLarp7 mediated rescue of pathogenic phenotype was seen in adult photoreceptors (Figure 3. A c, c’ vs b, b’). Acridine Orange staining of the larval eye-antennal discs revealed rescue in the *SCA8*(CTG112) associated cell death in the larval photoreceptor neurons (Figure 3. A c” vs b”). In addition, the axonal projections were visualized with anti-22C10 antibody immunostaining as it marks the developing neurons. Ectopic dLarp7 significantly improved the neuronal morphology and axonal projections (Figure 3. A c”’ vs b”’) which was impaired by pathogenic *SCA8(CTG112)* non-coding transcripts (Figure b”’). To assess the role of *spoon*, both hypomorphic allele *spoonbill^KG02745^* or overexpression allele *UAS-HA-spoonbill* were examined in combination with *dLarp7* and pathogenic *SCA8(CTG112)* lines. Interestingly, the rescue of *SCA8* associated pathogenic phenotype by dLarp7 was significantly reduced in combination with the hypomorphic allele, *spoonbill^KG02745^*. The pathogenic *SCA8* flies with dLarp7 overexpression along with reduced Spoon levels resulted in severe eye roughening (Figure 3. A d), with increase in number of fused ommatidia (Figure 3. A d’), elevated cell death (Figure 3. A d”) and disorganized neuronal morphology (Figure 3. A d”’). However, there was no additional rescue in the adult and larval neurodegeneration when both these proteins were co-expressed (Figure 3. A e, e’, e”, e”’). Quantification of adult photoreceptor phenotype indicated that depletion of Spoon interferes with dLarp7 mediated rescue of *SCA8* neuropathology (Figure 3. A’). Hence, the combinatorial function of both Spoonbill and dLarp7 is needed to mitigate *SCA8*(CTG112) associated pathogenicity. However, co-expression of both the proteins did not confer any additional protection to the neurons against pathogenic *SCA8*(CTG112) transcript induced neurodegeneration, possibly because the two proteins work together through a common pathway to alleviate the pathogenicity associated with *SCA8*(CTG112) transcripts.

Further the regulatory role of Spoon on dLarp7 levels were also examined. dLarp7 expression was reduced in Spoonbill depleted tissue as revealed by immunostaining and western blot (Figure 4. A, B). Western blotting against HA in lysates with *GMR-Gal4* driven *UAS-HA-dLarp7* in combination with hypomorphic *spoonbill^KG02745^,* revealed significant reduction in the levels of the dLarp7 (Figure 4. A). Depleted *spoon* resulted in no modulation of the expression of RNA binding protein *elav* (Figure 4. B b’ vs a’), however, dLarp7 levels were significantly reduced (Figure 4. B b vs a). To assess the physiological effect of depleting Spoon on dLarp7 functions, we next checked the level of the RNA target of dLarp7 protein. dLarp7 acts as a chaperone for *7SKsnRNA* (Nguyen et al. 2012). The level of *7SKsnRNA* was depleted in *spoonbill^KG02745^* when compared to that of the wild type control (Figure 4. C). Thus, we conclude that spoon positively regulates dLarp7 and this in turn control downstream targets of dLarp7 like *7SKsnRNA*.

**Figure 4.**
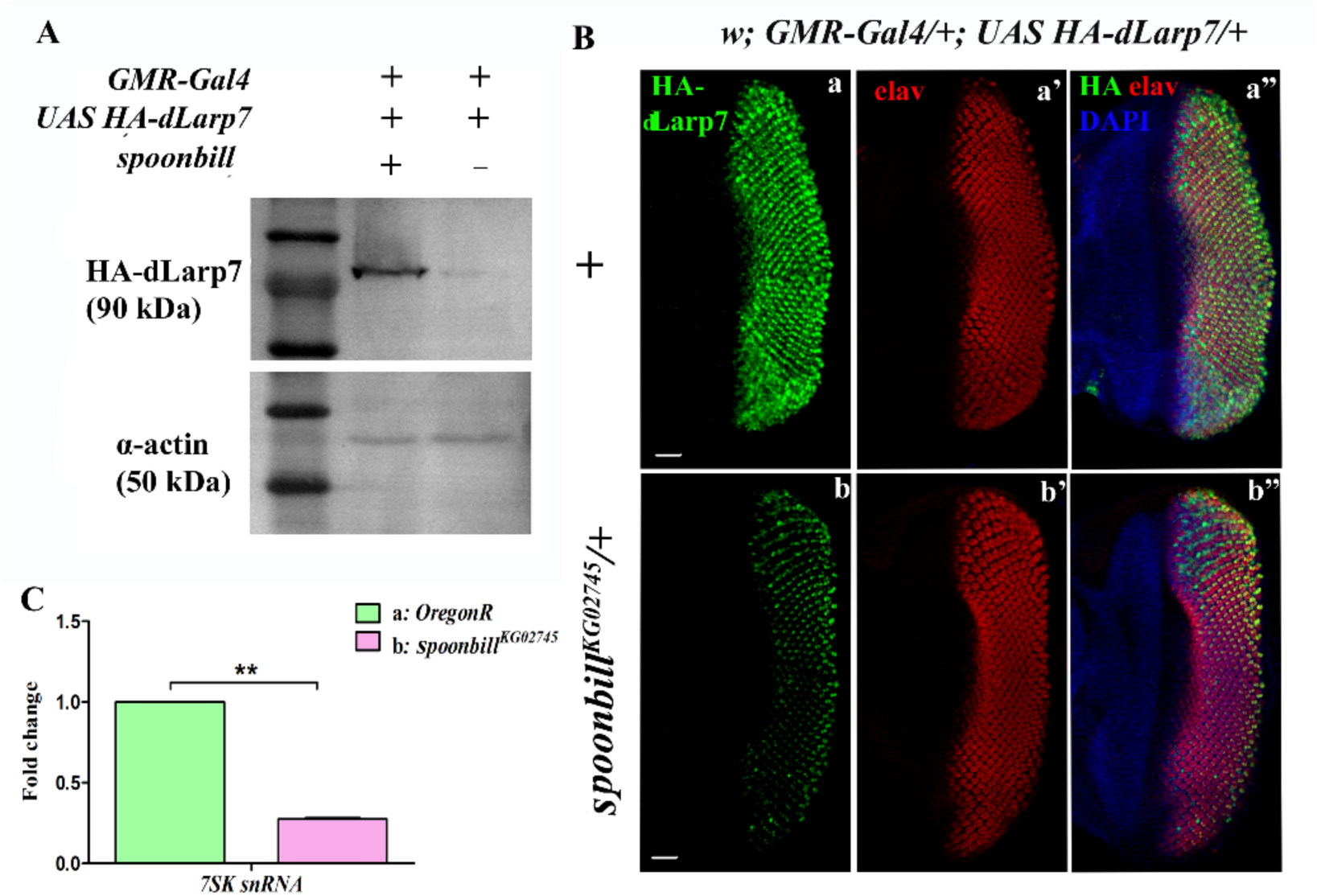
dLarp7 expression and function depends on availability of Spoonbill protein in *Drosophila* eye-antennal and adult brain. (A) Western blot by anti-HA antibody performed to detect the dLarp7 expression in lysates obtained from adult *Drosophila* heads, revealed depleted expression in *spoonbill^KG02745^, GMR-Gal4>UAS-HA-dLarp7* (upper panel) compared to *GMR-Gal4* driven *UAS-HA-dLarp7* alone (upper panel). *β-*Actin was used as input control for the lysates (lower panel). (B) Immunostaining against HA-dLarp7 showed depleted expression of dLarp7 in reduced availability of Spoonbill (b) compared to when physiological levels of Spoonbill is present (a). Counterstaining with anti-elav showed no variation in response to depletion of Spoonbill protein and marked organized photoreceptor neurons in both conditions (a’ and b’). The morphology of eye-antennal discs also showed no change in response to reduced Spoonbill expression which resulted in dLarp7 depletion (b” and a”). Scale bars, 50 µm. (C) Transcript level of *7SKsnRNA,* as seen by RT-PCR, depletes in *spoonbill^KG02745^*fly heads compared to wild type flies

### 4. dLarp7 depletes pathogenic *SCA8*(CTG112) transcripts and RNA foci formation

To gain an insight into the molecular mechanism of dLarp7-*SCA8*(CTG112) interaction, *in situ* hybridization was performed to detect pathogenic *SCA8*(CTG112) transcript status in the developing photoreceptor neurons. *GMR-Gal4* driven *UAS SCA8*(CTG112) formed large number of *SCA8* pathogenic RNA foci (Figure 5. A a, a’). The RNA foci are insoluble toxic aggregates of ribonucleoproteins formed by the trinucleotide expanded *SCA8*(CTG112) RNA which acquires secondary hairpin loops and sponges away vital RNA binding proteins (Mutsuddi et al., 2004). The number of toxic RNA foci formed by *SCA8*(CTG112) (Figure 5. A b, b’ vs a, a’) was dramatically reduced in the presence of ectopic dLarp7. Additionally, we also observed that majority of the foci size were smaller demonstrating marked suppression of pathogenicity mediated by *SCA8*(CTG112) transcripts. Hence, the suppression of *SCA8*(CTG112) associated pathogenic RNA foci by dLarp7 was also confirmed at the molecular level. In concordance with our genetic interaction studies, reduction of Spoonbill levels impeded suppression of pathogenic *SCA8* RNA foci formation by ectopic dLarp7 (Figure 5. A c, c’ vs a, a’). Overexpression of both, *spoon* as well as *dLarp7* resulted in no additional rescue of the toxic RNA foci formation in *Drosophila* eye-antennal discs than that by dLarp7 alone (Figure 5. A d, d’ vs a, a’). Quantitative analysis of the reduction in foci number with co-expression of dLarp7 and pathogenic *SCA8*, was highly significant (Figure 5. B). RT-PCR of *SCA8*(CTG112) transcripts further strengthened the above data and revealed a 0.5-fold reduction of *SCA8* transcripts in the presence of ectopic dLarp7 (Figure 5. C). Briefly, ectopic dLarp7 expression reduced the levels of pathogenic *SCA8* transcripts as well as the number and size of pathogenic RNA foci significantly. This reduction depended largely on the presence of Spoonbill as our study indicates that Spoon positively regulates dLarp7 expression and function. Thus, dLarp7 along with Spoon can act as a potent therapeutic endpoint for Spinocerebellar Ataxia 8 associated neurodegeneration.

**Figure 5.**
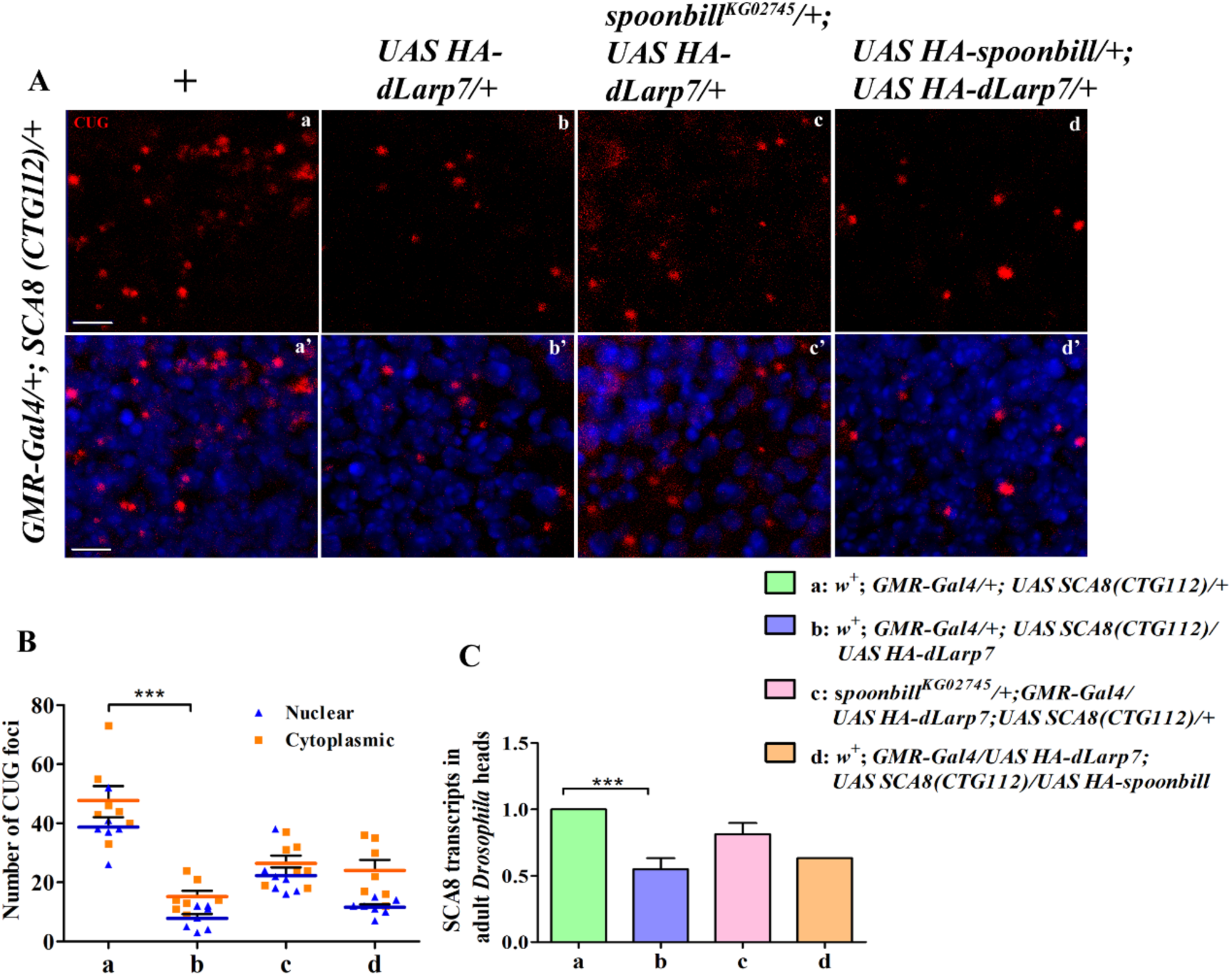
Pathogenic *SCA8* is suppressed by *UAS-HA-dLarp7*. (A) dLarp7 modulates *UAS SCA8(CTG112)* associated RNA foci formation in Spoon dependent manner. Fluorescence *in-situ* hybridization for *SCA8(CTG112)* specific riboprobes in *GMR-Gal4* driven *UAS SCA8(CTG112)* show formation of foci in cytoplasm and nuclei (a, a’). Overexpression of *dLarp7* results in massive reduction in the number and relative size of the foci formed (b, b’). Combining *spoonbill^KG02745^,* with *GMR-Gal4* driven *UAS SCA8(CTG112)* and *UAS-HA-dLarp7* results in increased number of foci (c, c’ vs b, b’), whereas co-expression of *UAS-HA-spoonbill,* in the same background results in no additional reduction in foci formation (d, d’ vs b, b’). (B) The number of foci were counted per unit area for an average 10 discs for each genetic combination. Overexpression of *dLarp7* results in drastic reduction in number of foci, from an average of nearly 50 to about 10 foci were seen in the larval eye antennal discs (b vs a). Combining *spoonbill^KG02745^,* with *GMR-Gal4* driven *UAS SCA8(CTG112)* and *UAS-HA-dLarp7,* the number of foci increases to an average of 25 (c vs b), whereas with co-expression of *UAS-HA-spoonbill,* in the same background results in formation of about 18 foci on an average (d, d’ vs b, b’). The reduction of number of foci formed is significant between b and a. To calculate statistical significance of the data Bonferroni pot test was added after two-way ANOVA analysis for the genetic interactions. Bars represent average ± SEM for multiple independent experiments. p values < 0.001***. Scale bars, 5 µm (A a, a’). (C) RT-PCR for *UAS SCA8(CTG112)* revealed about half a fold decrease in the *SCA8* transcripts (b vs a). This rescue was impaired as *spoonbill^KG02745^*, was brought in combination with *UAS-HA-dLarp7* in pathogenic *SCA8* background (c). Rescue in *SCA8* transcripts when *UAS-HA-spoonbill* was co-expressed with *UAS-HA-dLarp7* and *UAS SCA8(CTG112)*, is observed to be nearly similar to that by *UAS-HA-dLarp7* alone (d). The bars plotted represent average fold changes of two independent experiments ± SEM. To obtain the significance of the RT-PCR experiment, *t-*test was performed. p values < 0.001***

### 5. Pathogenic *SCA8*(CTG112) recruits dLarp7 protein into RNA-protein complex

Pathogenic RNAs with repeat expansions form toxic RNA foci, that recruit vital RNA Binding Proteins away from their physiological pool. We investigated whether dLarp7 is a part of the pathogenic *SCA8* RNA-protein complex that results in toxic RNA foci formation. RNA pull-down assay was done where *in-vitro* transcribed *SCA8*(CTG112) was added to the *GMR-Gal4* driven *UAS-HA-dLarp7* lysates. HA-tagged dLarp7 was pulled down and the interacting RNAs were reverse transcribed into cDNA. Interestingly, *SCA8* transcripts were pulled down only in samples with dLarp7 overexpression. No amplification was seen in samples with dLarp7 where Spoonbill was depleted (Figure 6. A). The level of dLarp7 in the protein lysates used for the pull-down assay was verified by western blotting against HA and Actin, respectively (Figure 6. A’). Our observations suggests that dLarp7 forms a biological complex with trinucleotide repeat expanded *SCA8*(CTG112) transcripts which is Spoonbill dependent. Amplification of *7SKsnRNA* was also obtained in samples with *UAS-HA-dLarp7* and *UAS-SCA8*(CTG112), reflecting the restoration of dLarp7 levels which then leads to stabilization of *7SKsnRNA* (Figure S4).

**Figure 6.**
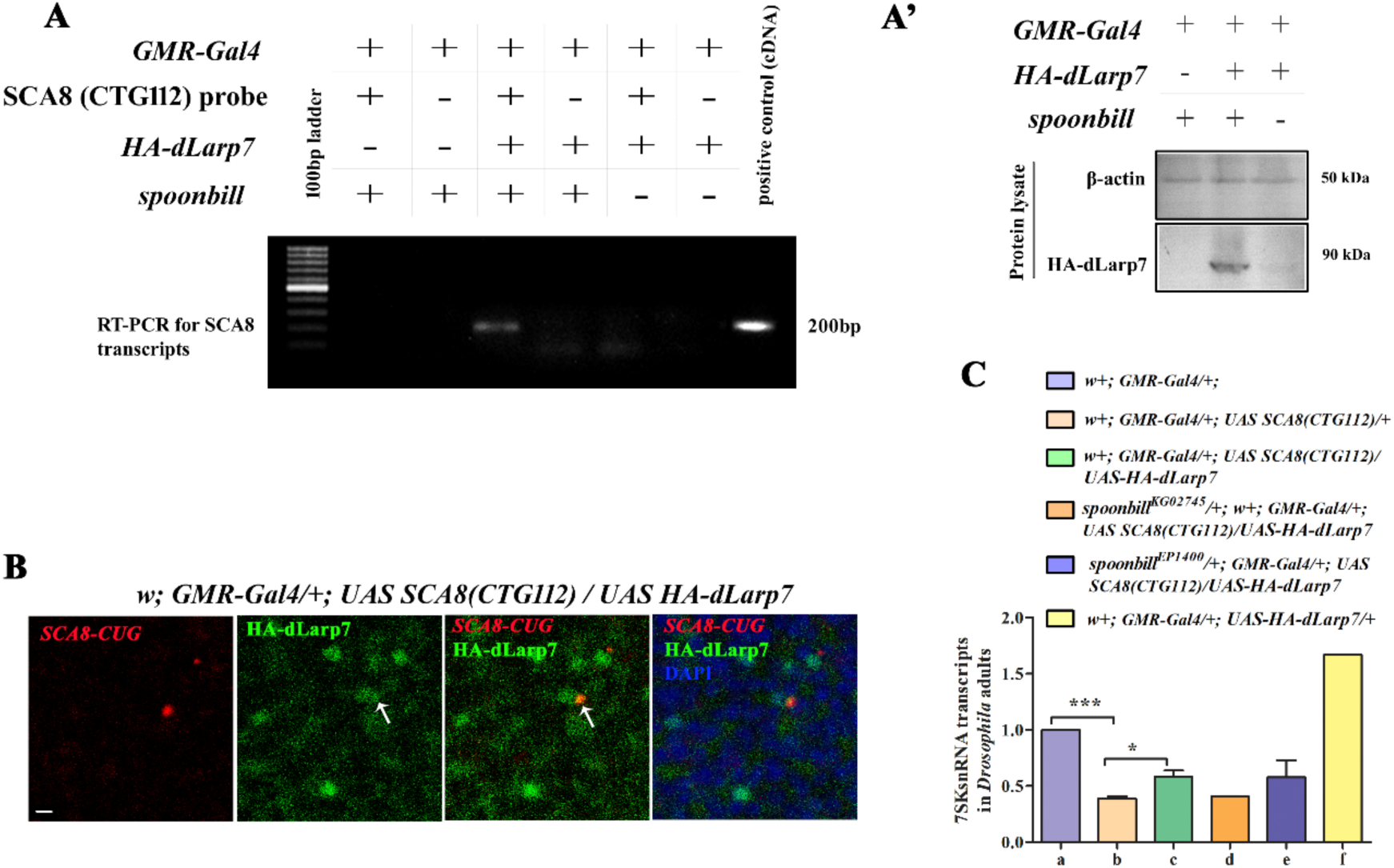
dLarp7 forms unusual complex with *UAS SCA8(CTG112)* in Spoonbill depended manner. (A) RT-PCR for *UAS SCA8(CTG112)* was performed using total cDNA as template, obtained from RNA co-immunoprecipitated with HA-dLarp7 when pulled down using HA-beads in adult *Drosophila* head lysates with different genetic combinations. Samples with *GMR-Gal4/+* was used as control for the experiment (Lane 2, 3). Samples with HA-dLarp7 were run in Lane 4, 5. Samples with HA-dLarp7 in Spoonbill depleted background were run in lanes 6, 7. *in-vitro* transcribed *SCA8*(CTG112) probes were added to the lysates for investigating physical association with the dLarp7 (Lane 2, 4, 6). In negative controls for all genetic combinations, *SCA8*(CTG112) probes were not added (Lane 3, 5, 7). cDNA of *GMR-Gal4* driven *UAS SCA8(CTG112)* was used as positive control for PCR. A single band at nearly 200bp was obtained in lane 4 in sample with *SCA8*(CTG112) probes added *in-vitro* to HA-dLarp7 lysates. RT-PCR from the total bound RNA immunoprecipitated using anti-HA conjugated beads reveals the presence of pathogenic *SCA8* in complex with HA-dLarp7 (Lane 4). Lane 1 – 100 bp DNA ladder. (A’) Western Blot for β-Actin (upper panel) and HA-dLarp7 (lower panel) was performed to check the input lysates used for the immunoprecipitation. (B) Immunofluorescence against dLarp7 coupled with RNA *in situ* hybridization reveals the recruitment of dLarp7 protein into *SCA8*(CTG112) foci. Scale bars, 5 µm. (C) RT-PCR for *7SKsnRNA* results in significant depletion in *UAS SCA8(CTG112)* overexpression compared to that from control *Drosophila* adult heads (b vs a). Overexpression of dLarp7 in *GMR-Gal4* driven *UAS SCA8(CTG112)* background results in partial but significant rescue in the *7SKsnRNA* transcript level (c vs b). Reduced expression of *spoon* in *GMR-Gal4* driven *UAS SCA8(CTG112)* and *UAS-HA-dLarp7* background results in depletion of *7SKsnRNA* transcript (d). Co-expression of *UAS-HA-spoonbill* and *UAS-HA-dLarp7* in the *GMR-Gal4* driven *UAS SCA8(CTG112)* rescues *7SKsnRNA* transcripts partially (e). RT-PCR for *7SKsnRNA* in *GMR-Gal4* driven *UAS-HA-dLarp7* was used as positive control for the experiment shows drastic upregulation of the transcripts (f). The bars plotted represent average fold changes of two independent experiments± SEM. To obtain the significance of the RT-PCR experiment, *t-*test was performed. p values < 0.001***, p values < 0.01*.

To visualize this RNA-protein interaction immunostaining coupled with FISH was performed. We observed colocalization of dLarp7 protein with *SCA8*(CTG112) RNA foci. Colocalization of dLarp7 protein and *SCA8*(CTG112) RNA foci was observed in spite of depletion in the number of the toxic RNA foci (Figure 6. B).

To assess the functional effect of this unusual RNA-protein association of *SCA8* and dLarp7, quantitative PCR for *7SKsnRNA* was performed as a read out of dLarp7 function. dLarp7 binds to the 3’ terminal U-rich stretch of the highly conserved *7SKsnRNA*, thereby stabilizing it for assembly into 7SKsnRNP (Markert et al. 2008). Interestingly, *7SKsnRNA* transcripts were depleted in pathogenic *SCA8*(CTG112) condition. Ectopic dLarp7 protein rescued the depletion of *7SKsnRNA* transcripts. Thus, it can be concluded that pathogenic *SCA8*(CTG112) RNA foci recruit the dLarp7 proteins along with other RBPs like Spoonbill. This depletes the RBP from its physiological pool, resulting in loss of the target RNA like *7SKsnRNA* (Figure 6. C).

Briefly, dLarp7, has been identified as a novel interacting partner of Spoonbill protein, which suppresses pathogenic *SCA8*(CTG112) neurodegenerative phenotypes in a *spoon* dependent manner. At the molecular level dLarp7 depleted the toxic RNA foci along with the reduction of trinucleotide repeat containing pathogenic transcripts of *SCA8*(CTG112). The recruitment of dLarp7 into the *SCA8*(CTG112) RNA foci results in the depletion of dLarp7 from its physiological pool and this in turn leads to degradation of its target, *7SKsnRNA*. We put forward a novel hypothesis that perhaps, loss of *7SKsnRNA* disturbs the cellular balance between splicing and transcription and might be one of the vital molecular mechanisms that contributes to *SCA8*(CTG112) associated neurodegeneration. While ectopic expression of dLarp7 rescues the *SCA8*(CTG112) associated neurodegeneration, as the physiological pool of dLarp7 and *7SKsnRNA* is partially restored.

### 6. SMN protein, suppresses Spinocerebellar Ataxia 8 associated neurodegeneration

We furthered our investigation to assess the role of interacting partners of dLarp7 in the disease pathogenicity of *SCA8*(CTG112). We evaluated the interaction between dLarp7 and SMN protein, whose recessive mutation causes neurodegenerative spinomuscular atrophy. Immunoprecipitation was done using α-FLAG to pull-down FLAG-SMN protein followed by immunoblotting against dLarp7. It was observed that the two proteins form complex in the photoreceptor neurons of *Drosophila* (Figure 7. A). Co-immunostaining using α-dLarp7 and α-HA (Figure 8. B) to visualize dLarp7 and α-FLAG for SMN protein revealed partial colocalization in the photoreceptor neurons of *Drosophila* larvae (Figure 7. B c’, g’).

**Figure 7.**
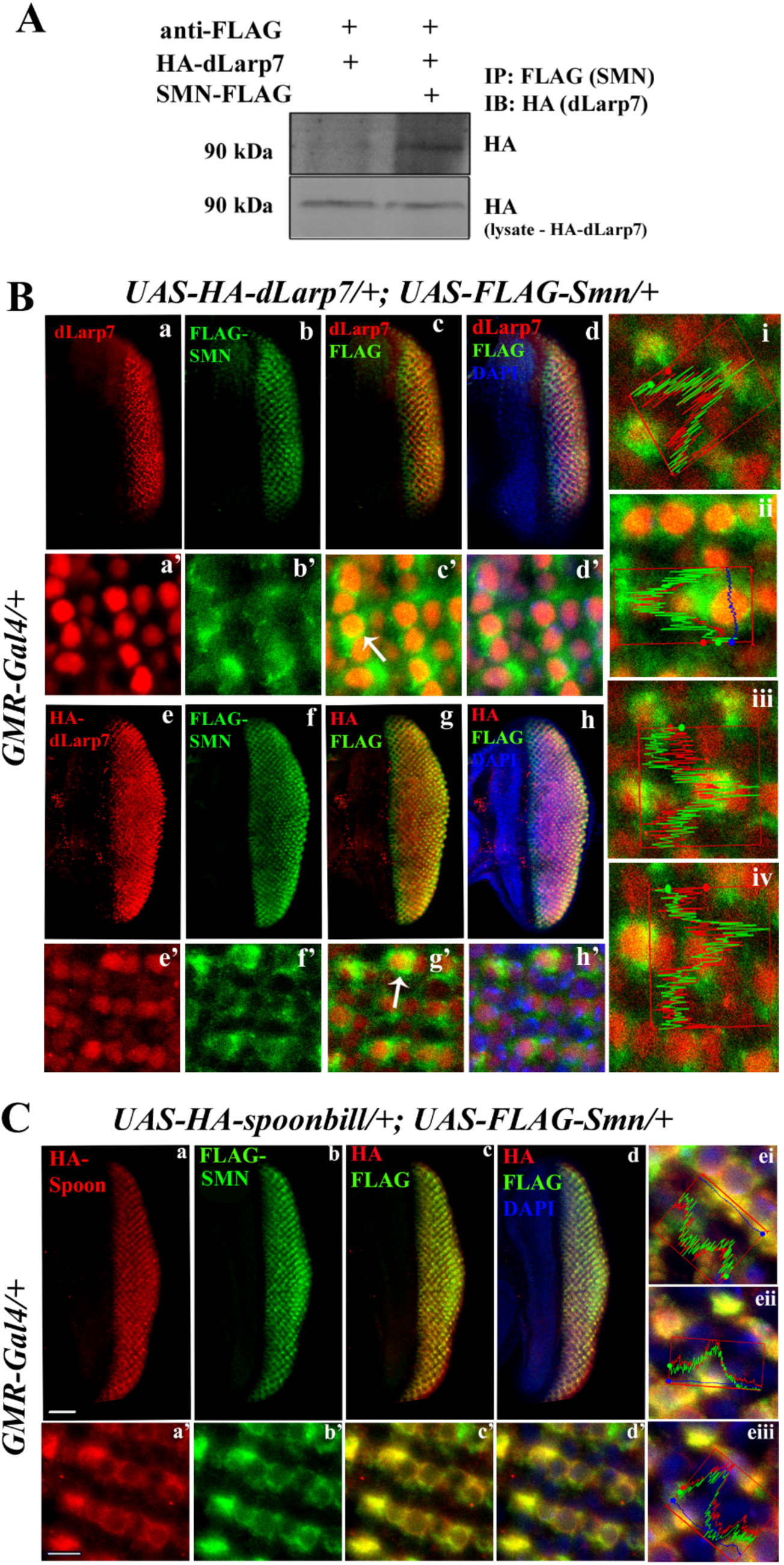
SMN protein colocalizes with dLarp7 and Spoonbill in *Drosophila* photoreceptors. (A) (B) Colocalization of SMN and dLarp7 proteins in the eye-antennal discs. *UAS-FLAG-Smn* is co-expressed with *UAS-HA-dLarp7* in the *GMR-Gal4* domain. dLarp7 is marked with anti-dLarp7 (a, a’) or anti-HA (e, e’) and SMN is marked with anti-FLAG (b, b’, f, f’). merged channel shows partial co-localization of the two proteins as yellow spots (white arrows in c’, g’). Intensity profiles show overlapping peaks for SMN and dLarp7 (i-iv). (C) Colocalization of Spoonbill and dLarp7 proteins in the eye-antennal discs. Spoonbill was marked with anti-Spoon antibody (a, a’). Merged channel show colocalization of the two proteins to a large extant (c’). Intensity peaks validate overlapping signals from both channels indicating a significant colocalization of the two proteins (i-iv). Scale bars, 50µm (a), 5 µm (a’).

**Figure 8.**
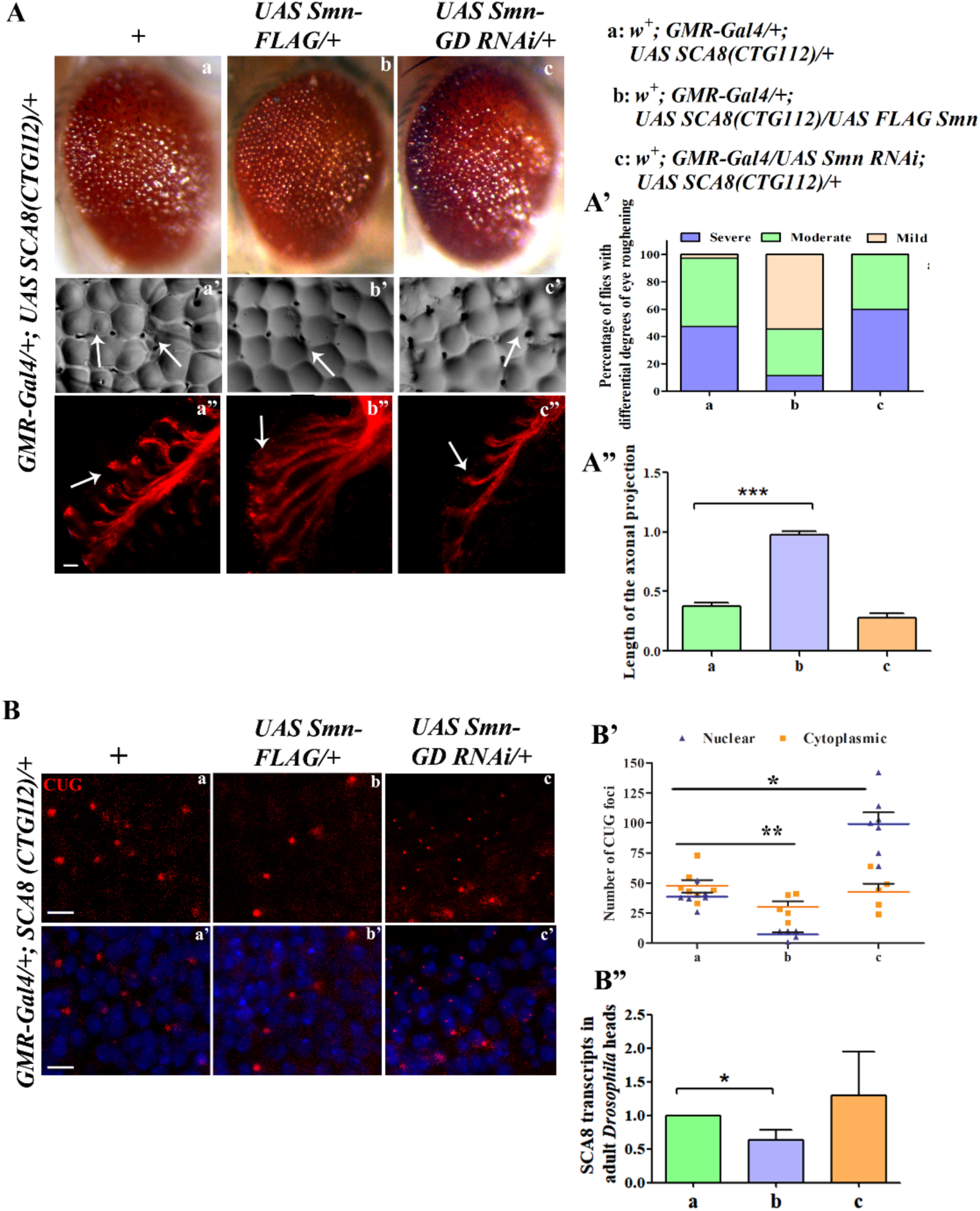
SMN modulates *UAS SCA8(CTG112)* molecular pathogenesis. (A) *UAS SCA8(CTG112)* associated eye roughening (a), fused disorganized ommatidia (a’) and degenerated axonal projections (a”) is rescued when combined with *UAS-FLAG-Smn* (b, b’, b”) in the *GMR-Gal4* domain. Upon combining *UAS-Smn RNAi* with the *UAS SCA8(CTG112)*, the neurodegenerative phenotype is further deteriorated (c, c’, c”). (A, A’) Quantification of the adult readouts and axonal projections indicate pronounced regulation of *SCA8* associated neurodegeneration by SMN protein. (B) Smn modulates the *UAS SCA8(CTG112)* associated foci formation. Fluorescence *in-situ* hybridization for *SCA8*(CTG112) specific riboprobes in *GMR-Gal4* driven *UAS SCA8(CTG112)* results formation of foci in cytoplasm and nuclei (a, a’). Overexpression of *Smn* results in reduction in the number of the foci (b, b’). Depleting *Smn* expression by combining *UAS-Smn RNAi,* with *GMR-Gal4* driven *UAS SCA8(CTG112)* results in dramatic increase in the number of nuclear RNA foci (c, c’ vs a, a’). (A’) The number of RNAfoci were counted per unit area in eye-antennal discs for each genetic combination. The modulation in RNAfoci number in response to varied doses of SMN was found to be significant (b, c vs a). To calculate statistical significance of the data Bonferroni pot test was added after two-way ANOVA analysis for the genetic interactions. (B”) *SCA8(CTG112)* transcripts are depleted significantly when *Smn* is overexpressed in adult heads. Bars represent average ± SEM for two independent experiments. p values < 0.001***. Scale bars, 5 µm (A a, a’).

Interaction with Spoonbill and SMN, was also assessed using α-Spoon and α-FLAG (Figure 7. C). Surprisingly, the immunostaining pattern for both the proteins in the photoreceptors of *Drosophila* was seen to be largely similar, indicating a significant colocalization of Spoon and SMN proteins (Figure 7. C c’, d’).

Following establishment of the novel interaction of Spoon, dLarp7 with SMN proteins in *Drosophila* photoreceptors and brain, we next assessed the role of *Smn* in Spinocerebellar Ataxia 8 associated neurodegeneration model. Interestingly, overexpression of *Smn* in pathogenic *SCA8*(CTG112) background, suppressed the neurodegenerative phenotype significantly (Figure 8. A b, b’ vs a, a’). In the third instar larval stage the manifestation of expanded *SCA8*(CTG112) can be depicted by degenerated axonal projections. Overexpression of *Smn* resulted in massive rescue of the axonal projections (Figure 8. A b” vs a”). On similar lines, *Smn* depletion led to enhancement of the pathogenic *SCA8*(CTG112) phenotype in the larval and adult photoreceptors (Figure 8. A c-c” vs a-a”). This indicates that Smn protein contributes to *SCA8*(CTG112) associated neurodegeneration. No change in the *UAS SCA8(CTG9)* induced rough eye phenotype was seen on SMN modulation (Figure S5).

This observation also correlated with the assessment at the molecular level. Overexpression of *Smn* depleted the expanded *SCA8*(CTG112) transcripts (Figure 8. B”). Fluorescence *in-situ* hybridization (Figure 9 B) against expanded pathogenic *SCA8*(CTG112) revealed significant reduction in nuclear RNA foci with *Smn* overexpression. Interestingly, depleting SMN protein in pathogenic *SCA8*(CTG112) background resulted in formation of more abundant but smaller RNA foci which were largely nuclear in localization (Figure 8. B, B’). This data clearly suggests that SMN protein suppresses the *SCA8*(CTG112) associated neurodegeneration by depleting the pathogenic *SCA8*(CTG112) transcripts and toxic RNA foci. Depleting SMN protein in the *SCA8*(CTG112) background, led to increased number of toxic nuclear RNA foci which directly correlates with the disease severity. Hence, we have identified *Drosophila* homolog of SMN protein, implicated in SMA, as a novel modulator of *SCA8*(CTG112) associated neurodegeneration.

**Figure 9.**
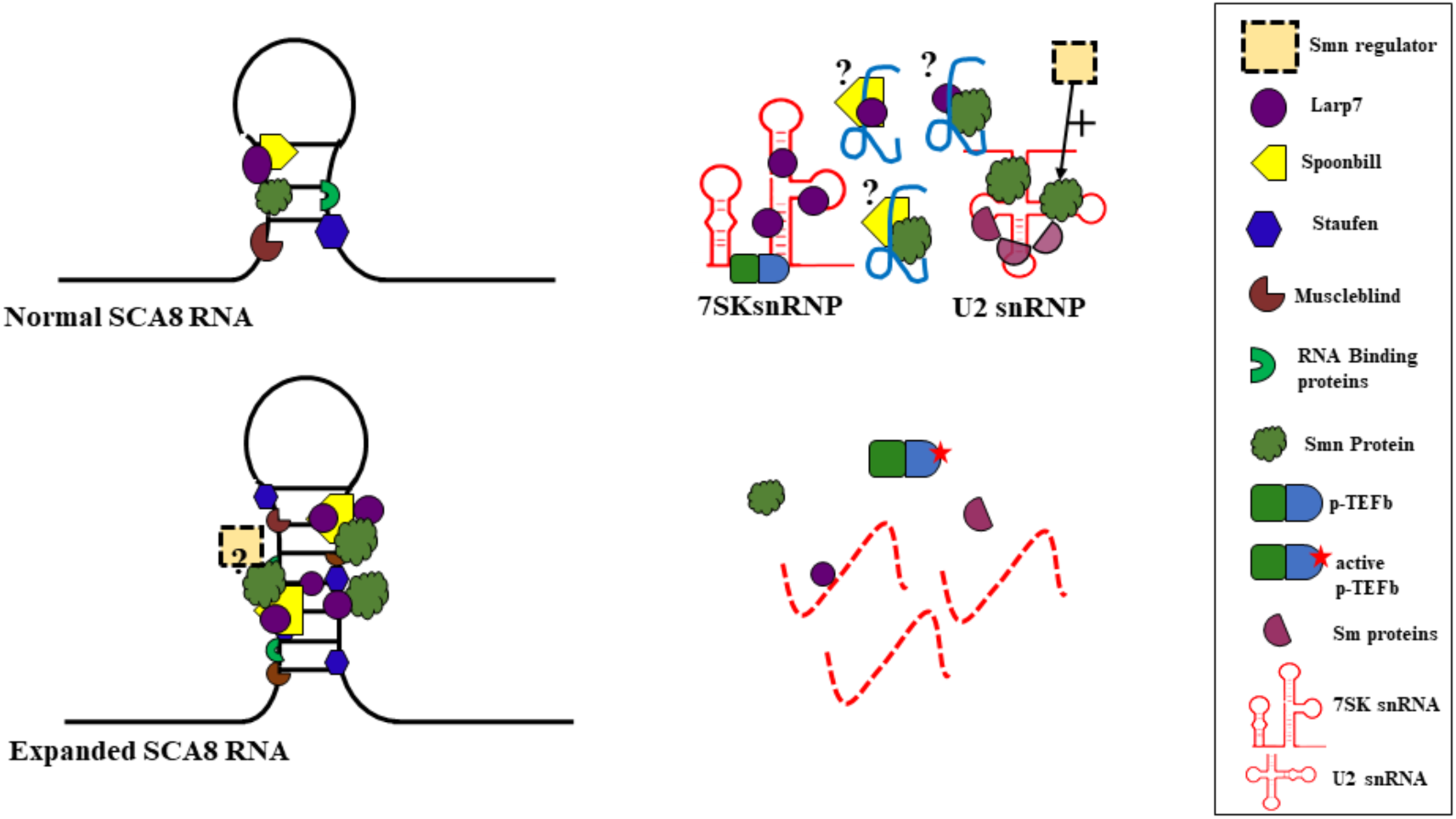
Hypothetical model of neurons with normal *SCA8* transcripts vs neurons with pathogenic *SCA8* transcripts. The RNA Binding proteins like Spoonbill, dLarp7, SMN maintain cellular RNA metabolism. In the presence trinucleotide repeat expanded *SCA8* RNA, vital RBPs like Spoonbill get recruited, that in turn engages other interacting partners like dLarp7 and SMN into the pathogenic *SCA8* RNA foci. This results in degradation of RNA targets of these critical proteins. Imbalance of splicing and transcription depending on these vital RBPs perhaps contribute to the expanded *SCA8* associated neurodegeneration. Increasing the dose of either of these proteins restore the physiological pool thereby reducing the *SCA8*(CTG112) associated neurodegeneration.

## DISCUSSION

*Drosophila* models of neurodegeneration have contributed immensely to our understanding of the disease pathogenesis. The fly model of Spinocerebellar ataxia 8 demonstrated cardinal features of the disease that facilitated screening for novel modulators, like Spoonbill, that are found to be vital for neuronal functions and development (Mutsuddi, et al., 2004; Mutsuddi, et al., 2005; Tripathi et al., 2016; Tripathi et al., 2017; Das et al. 2019; Das et al. 2023). The *Drosophila SCA8* model has helped in establishing the pathomechanisms of recruitment of vital RBPs into toxic RNA foci. The spatio-temporal dynamics of the RNA foci size and components are of pivotal importance in unraveling the molecular machinery underlying the disease as well as developing therapeutic endpoints.

In this pursuit we have employed a proteomic approach to identify interacting partners of the established modulators of the *SCA8* associated neurodegeneration in *Drosophila* disease model. In the current study we have established the role of an RBP dLarp7, a novel interacting partner of Spoon, in *SCA8 Drosophila* model. Using multiple molecular genetic tools, we have established that dLarp7 is recruited into the pathogenic *SCA8* RNA foci, thus overexpression of dLarp7 suppresses *SCA8* associated neurodegeneration by restoring its physiological pool of the RBP. Using protein interaction approach, we have also identified *Drosophila* SMN as a novel interacting partner of dLarp7 and Spoon. Homozygous recessive mutation of human *SMN1* leads to a neurodegenerative disease SMA/spinomuscular atrophy, where degeneration of the motor neurons are observed (MIM 253300). Interestingly, SMN also modulate *SCA8* associated molecular pathology and neurodegeneration. Our study identified two RBPs that are not just critical for cellular functions but their human homologs are strongly associated with neurodevelopmental disorders. This suggests occurrence of shared molecular machinery underlying such diseases. Perhaps, reduced functions of such RBPs either due to loss of function mutation or due to unusual recruitment into RNA foci renders (Figure 9.) the neurons susceptible to degeneration, as seen in multiple neurodevelopmental disorders.

*Drosophila* model of *SCA8*, like other disease models, have identified critical factors that contribute in non-coding RNA associated neurodegeneration. Genetic screen employing *Drosophila* model of *SCA8* identified *muscleblind* as a suppressor of neurodegeneration (Mutsuddi et al., 2004). As a proof of principle its mammalian orthologue MBNL1 was found to co-localize with the *SCA8* RNA foci in human cerebellar autopsy and mice brain cells (Daughters et al., 2009).

Interestingly, MBNL1 was also identified to be recruited into RNA foci associated with a neuromuscular disorder Myotonic Dystrophy 1 (DM1). Expansion of CTG repeats within the 3′ untranslated region of the DMPK gene in chromosome 19q13.3 results in DM1 (Aslanidis et al., 1992; Mahadevan et al., 1992; Mahadevan et al., 1993; Tiscornia & Mahadevan, 2000). The two disorders myotonic dystrophy1 and spinocerebellar ataxia 8 share a similar genetic etiology i.e., expansion of CTG repeats in the untranslated region of specific genes. Similar to pathogenic *SCA8* RNA, patients with DM1 have ribonuclear foci formed in patient fibroblasts and muscle biopsies (Mankodi et al., 2001; Pettersson et al., 2015; Ranum & Day, 2004). Sequestration of MBNL by the expanded transcripts, have been correlated with abnormal transcript processing and splicing abnormalities in numerous genes including *Troponin T* that contributes to the development of muscle weakness and myotonia (Nakamori et al., 2007). Staufen is another modifier of *SCA8* phenotype in *Drosophila*. Its mammalian homolog Stau1 rescues aberrant splicing profile of pre-mRNAs in DM1 (Bondy-Chorney, Crawford Parks, et al., 2016). Additionally, the suppression of DM1 phenotype by overexpression of Spoon and dLarp7 was also observed using the *Drosophila* model (unpublished).

Shared molecular pathology has also been shown in motor neuron diseases SMA and ALS, with distinct genetic etiology. Clinically, both are found to co-occur within families (Corcia et al. 2018). Accumulating evidence indicates intensive molecular overlap between the two diseases. *Smn* overexpression, in *Drosophila* cortical neurons alike, suppresses ALS-linked FUS mutation induced neuronal degeneration (Casci et al., 2019). Model based studies have identified functional relationship between Gemin3, a core component of SMN complex involved in SMA, and TDP-43 and FUS, which are associated with ALS (Cacciottolo et al., 2019). Motor neurons derived from SMA and ALS patients with reduced levels of *SMN1* have been shown to be more susceptible to cell death. Similarly, increasing doses of Smn was found to be beneficial for SOD1 and TDP-43 mutant mice models (Perera et al., 2016). Additionally, DM1 linked Muscleblind may also be involved in ALS as, it can regulate mislocalization of the SMN protein in (ALS)-linked FUS models of mice cortical neurons and *Drosophila* (Casci et al., 2019)

Our study has identified a novel functional association of *Drosophila* SMN protein in Spinocerebellar Ataxia 8 associated neurodegeneration. Additionally, we have demonstrated that the SMN protein interacts with Spoonbill and dLarp7 proteins that have been identified as suppressors of *SCA8* associated neurodegeneration. Based on our observations, SMN is a potent suppressor of pathogenic *SCA8* neurodegeneration. We hypothesise that the three proteins work together through overlapping or independent pathways in maintaining proper neuronal function. Their suppressive role in *SCA8* associated neurodegeneration opens up multiple therapeutic prospects for this neurodegenerative disorder. Deeper investigations into the role played by the RNA targets of Larp7 and SMN in neurodegenerative disorders and functional relevance of their interaction in neurons can help unravel shared molecular modulations for a range of neurodegenerative disorders. Presence of human homologs and orthologs for dLarp7 and SMN and their association with other neurodevelopmental disorders open up broader prospectives of these candidates as potent therapeutic targets for alleviating diseases. Identification of shared molecular mechanisms across a wide range of neurodegenerative disorders can help in early diagnosis and therapeutic management. Some vital RNA Binding Proteins that facilitate critical RNA functions in neurons share pathogenic pathways which may underlie multiple neurodegenerative disorders.

Investigations into cross-disease correlations can unravel the shared therapeutic endpoints. Interestingly, *SCA8* patients demonstrate ataxic phenotypes that are similar to ALS. Hence *SCA8* mutations can also be associated with dysfunctions of neurons outside the cerebellum (Kim et al., 2013), the site for the pathogenicity of expanded *SCA8*. Studying overlapping mechanisms between neurodegenerative disorders can reveal novel therapeutic targets that are independent of the site of disease affecting stabilization of neuronal homeostasis, repair and thus reinforcing the neuromuscular junctions (Comley et al., 2016; Wareham et al., 2022).

## Author contributions

Conceptualization, R.D and M.M.; Methodology, R.D., B.T. P.P. and B.M; Investigation, R.D., A.M. and M.M; Software and Formal Analysis, R.D. and MM; Writing Original Draft, R.D. and M.M; Supervision, M.M.

## Financial declarations

We do not have funds for covering OAP charges.

## Acknowledgments

We acknowledge the Bloomington Drosophila Stock Centre and Vienna Drosophila Resource Centre for fly stocks. We thank Prof. David Price, Dr. Baskar Bakthavachalu and Prof. Udai Bhan Pandey, Bloomington Stock Centre, DGRC and DSHB for providing us with the fly stocks, plasmid and antibodies used in this study. We acknowledge the real time facility at ISLS (Interdisciplinary Centre for Life Sciences) and Department of Molecular and Human Genetics. We also acknowledge confocal facility at ISLS and Banaras Hindu University. R.D., P.P., B.T. and B.M. were supported by fellowship from the Council of Scientific and Industrial Research, Government of India. This work was funded by CRG/2022/001003 from ANRF and IoE incentive grant from Banaras Hindu University to MM.

## Supplementary Information

**Fig S1:**
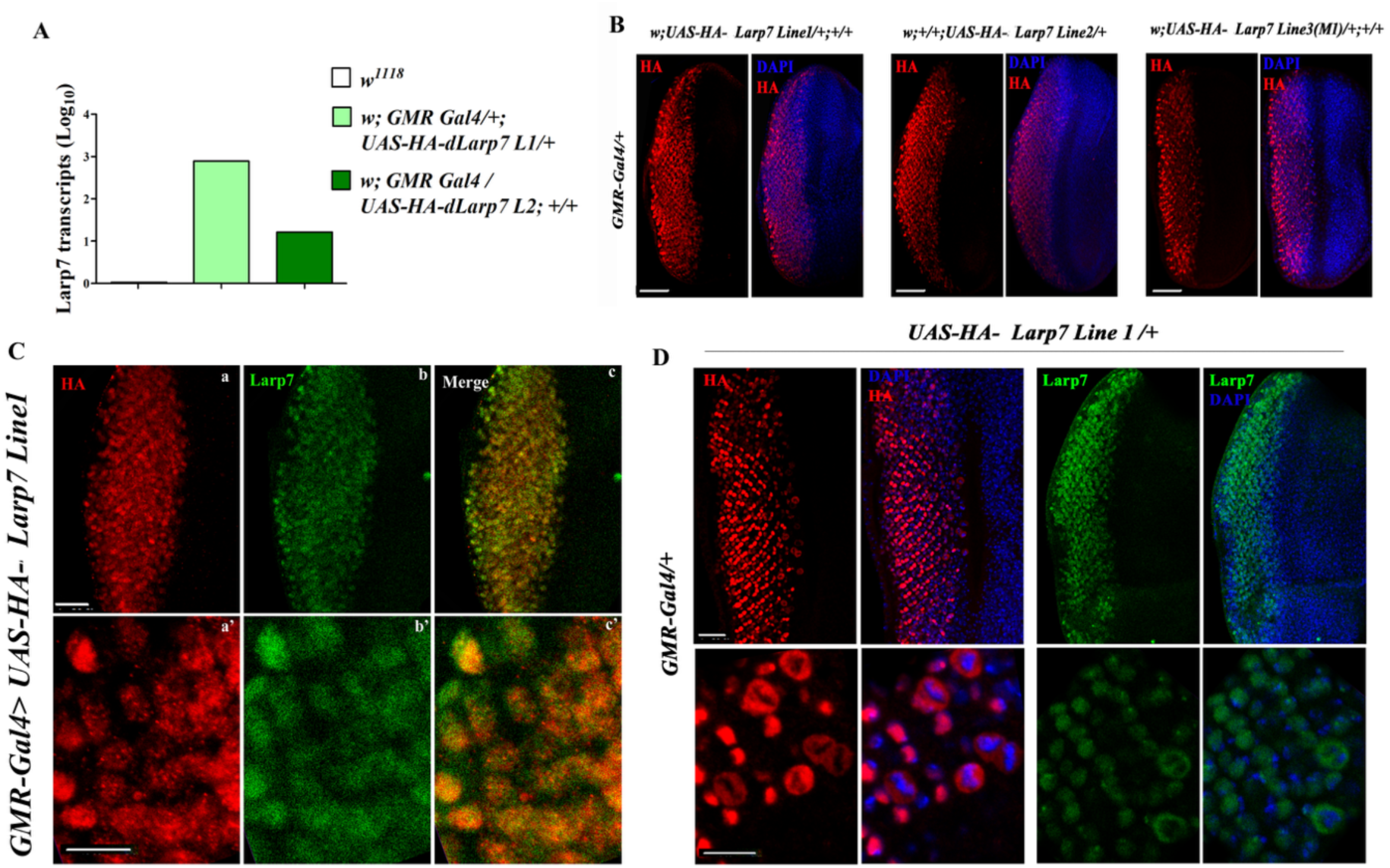
Validation of transgenic *Drosophila* lines with overexpression HA tagged dLarp7. Briefly, the complete coding sequence of dLarp7 *UAS-HA-dLarp7* lines were generated with reference genome FB2016_05. The 1.8kb CDS was cloned into *pUAST* vector between *KpnI* and *XbaI* with HA tag added to the 5’ end of the CDS. The recombinant plasmid was isolated using QIAGEN midi-prep kit and sequence verified. The plasmid was microinjected into the pole cells of the *w^1118^*embryo employing standard procedures for embryonic germ line transformation. Screening for the positive transformants was done by mini-white expression in fly eyes, that lead to identification of two lines. The chromosomes with the transgenes in the three transformant lines, were mapped for further studies. Various experiments were performed to validate the lines by expressing the transgenes using the UAS-Gal4 system (Brand & Perrimon, n.d.).Transgenic cDNA were expressed in the ommatidial region of the eye-antennal discs using *GMR-Gal4* driver. **(A)** RT-PCR was performed for dLarp7 using the RNA extracted from *Drosophila* heads of *GMR-Gal4* driven *UAS-HA-dLarp7* lines. Log fold change showed a very significant upregulation of dLarp7 in the lines. This proves that the overexpression lines result in upregulation of dLarp7. **(B)** Immunostaining against anti-HA revealed specific staining in the *GMR-*domain for the lines. Based on immunostaining Line3 was not used further. **(C)** Co-immunostaining against anti-HA and anti-dLarp7 (Nguyen et al., 2012) show complete overlap of signals which validated that the lines which expresses HA-tagged dLarp7. **(D)** While dLarp7 was seen to be largely nuclear, transient localization of dLarp7 in the cytoplasm in the region close to morphogenetic furrow was seen using anti-HA and anti-dLarp7 antibodies.

**Figure S2.**
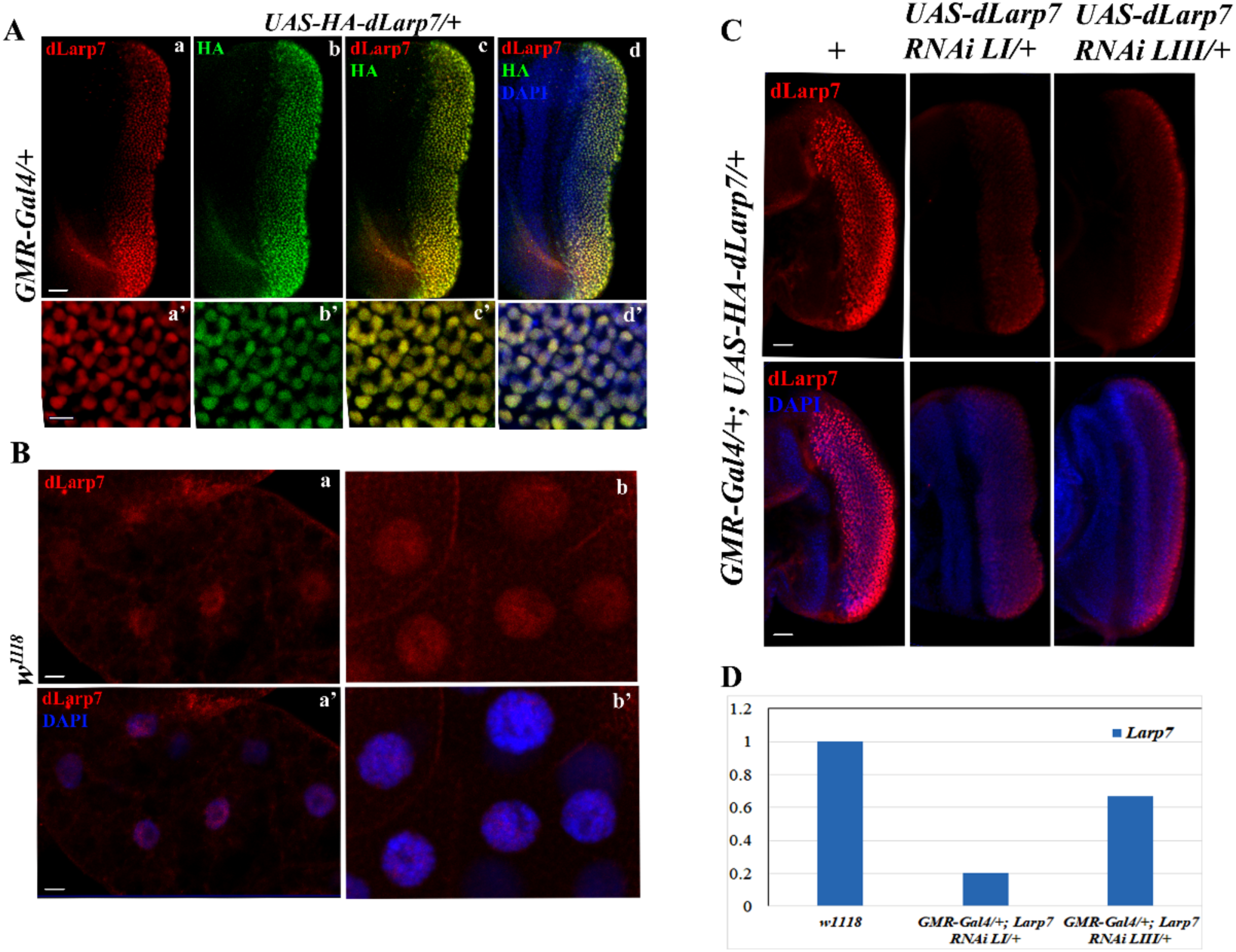
Verification of anti-dLarp7 antibody and transgenic *UAS-dLarp7 RNAi* fly lines. (A)Antibody against dLarp7 marked the protein in the photoreceptors of the eye antennal discs of in *GMR-Gal4* driven *UAS-HA-dLarp7*. (B) Anti-dLarp7 can detect the nuclear protein in the nucleus of salivary glands and fat tissues of *w^1118^*. (C) Immunostaining against dLarp7 showed depletion of dLarp7 in *GMR-Gal4* driven *UAS-dLarp7 RNAiLI* and *UAS-dLarp7RNAiLIII*. (D) RT-PCR against *dLarp7* show depletion of the target transcripts in the RNAi lines compared to the *w^1118^* control.

**Figure S3.**
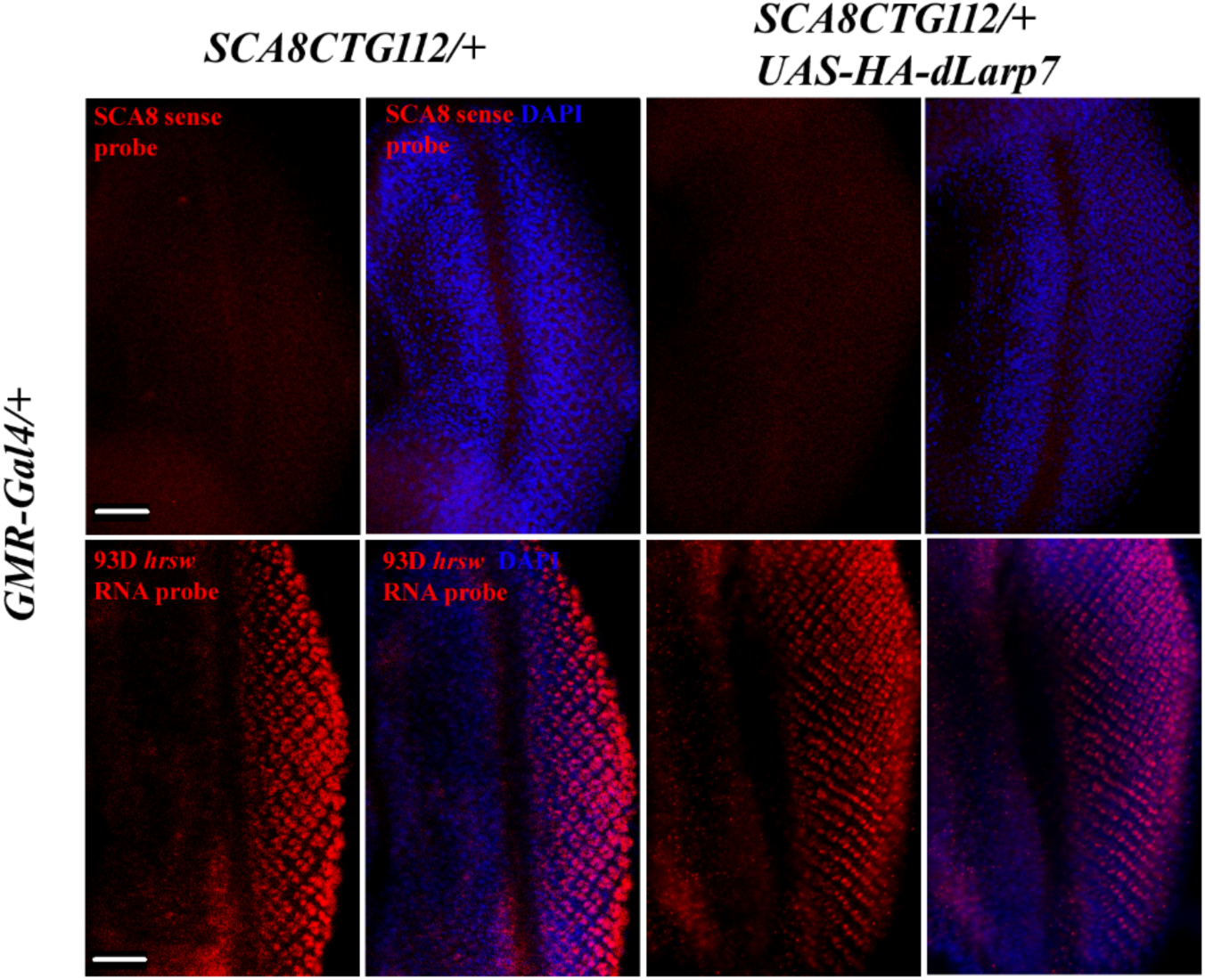
Controls for *in situ* hybridization. Positive control *hsrw 93D* (lower panel) and negative control *SCA8* sense probe (upper panel) show no difference between SCA8(CTG112) and *UAS-HA-dLarp7* combined with *UAS-SCA8(CTG112)*.

**Figure S4.**
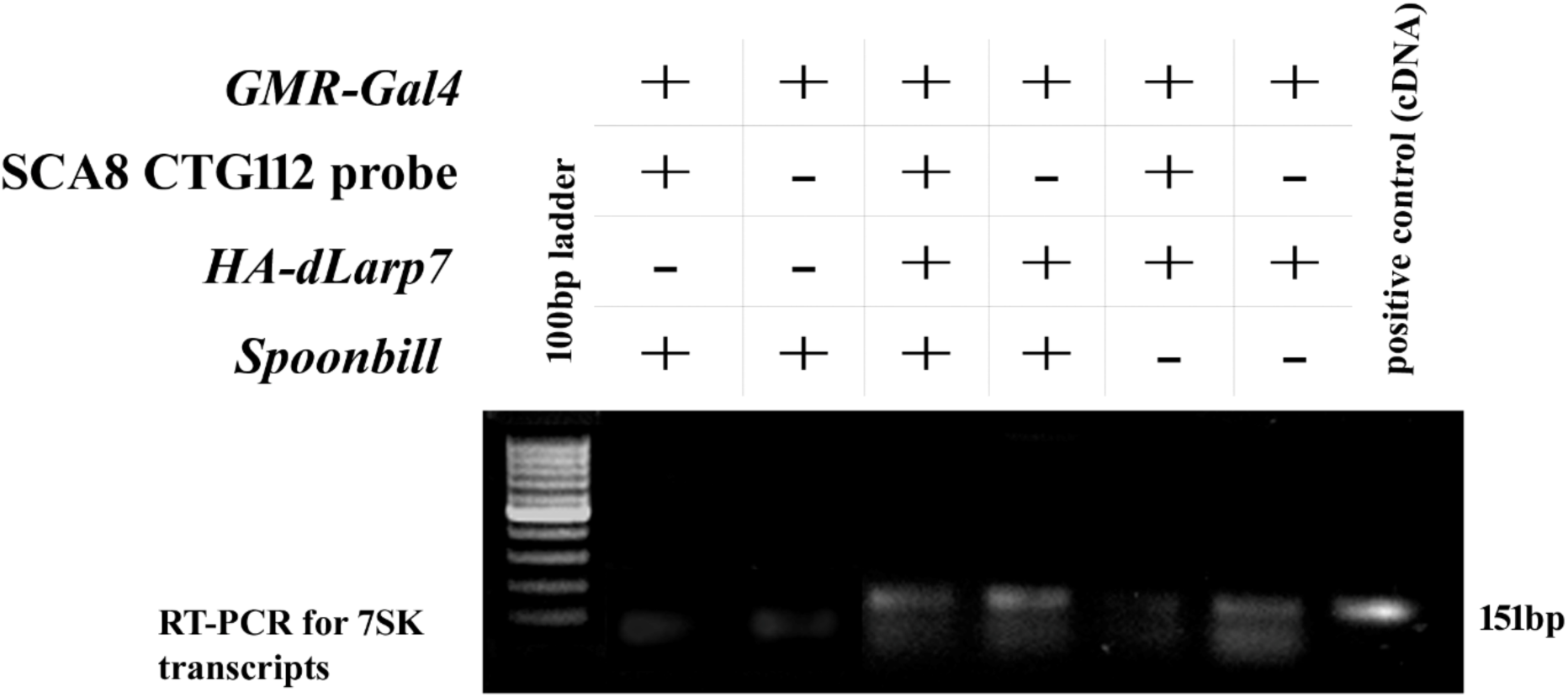
Internal control for RNA-immunoprecipitation. RT-PCR against *7SKsnRNA* of the cDNA generated from the pulled-down RNA reveals the presence of the transcript in all genetic combination. The band is relatively faint in genetic combination with depleted *spoon,* since Spoonbill positively regulates dLarp7 expression.

**Figure S5.**
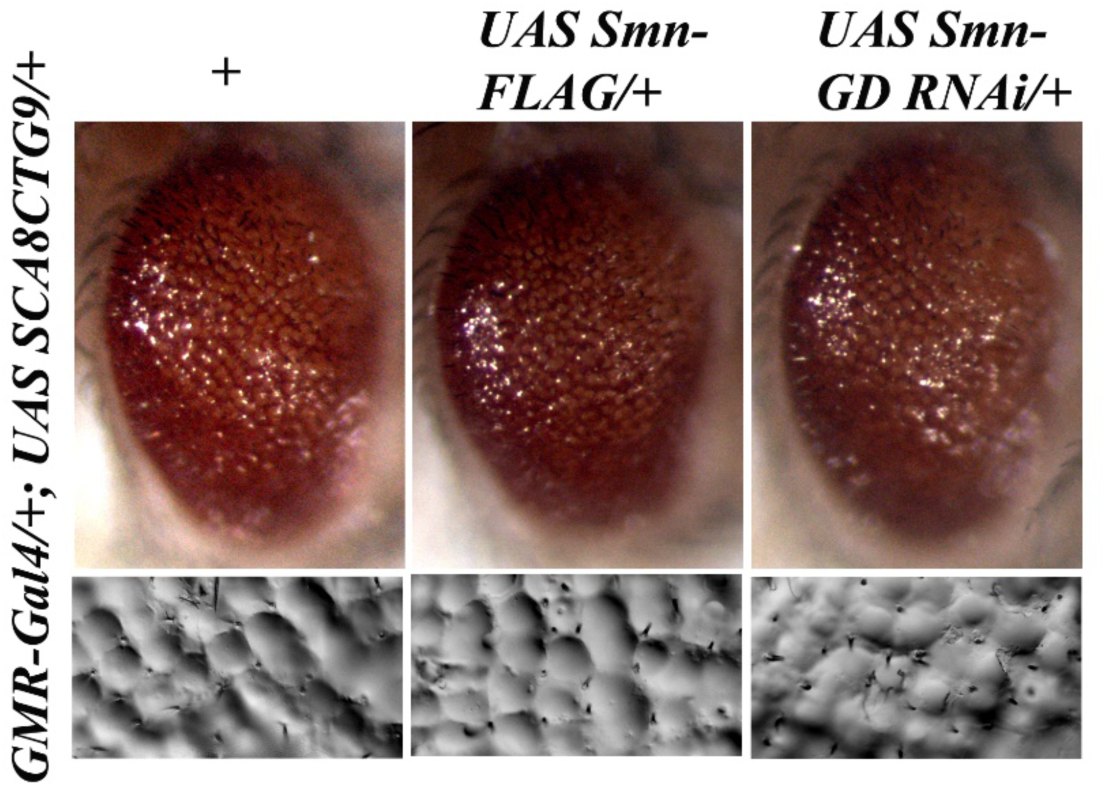
Modulation of Smn do not alter non-pathogenic SCA8(CTG9) phenotype. Overexpression or loss of function of Smn do not alter non-pathogenic SCA8(CTG9) associated eye roughening.

